# Diurnal variation in skeletal muscle mitochondrial function dictates time of day-dependent differences in exercise capacity

**DOI:** 10.1101/2024.09.02.610727

**Authors:** Subhash Khatri, Souparno Das, Anshit Singh, A Shabbir, Mohit Kashiv, Sunil Laxman, Ullas Kolthur-Seetharam

## Abstract

Exercise impinges on almost all physiological processes at an organismal level and is a potent intervention to treat various diseases. Exercise performance is well established to display diurnal rhythm, peaking during the late active phase. However, the underlying molecular/metabolic factors and mitochondrial energetics that possibly dictate time-of-day exercise capacity remain unknown. Here, we have unraveled the importance of diurnal variation in mitochondrial functions as a determinant of skeletal muscle exercise performance. Our results show that exercise-induced muscle metabolome and energetics are distinct at ZT3 and ZT15. Importantly, we have elucidated key diurnal differences in mitochondrial functions that are well correlated with disparate time-of-day dependent exercise capacity. Providing causal evidence, we illustrate that loss of Sirtuin4 (SIRT4), a well-known mitochondrial regulator, abrogates diurnal variation in mitochondrial functions and consequently abolishes time-of-day dependent exercise performance. Therefore, our findings unequivocally demonstrate the pivotal role of baseline skeletal-muscle mitochondrial functions in dictating diurnal exercise capacity.

## INTRODUCTION

Circadian rhythms orchestrate nearly all physiological processes in higher eukaryotes, including exercise, by integrating metabolic inputs with time-of-day signals (1). Emerging research indicates that muscle physiology and exercise capacity are strongly influenced by circadian rhythms, both in rodents (2, 3) and humans (4). Muscle fatigue, a critical determinant of exercise performance, is influenced by both central fatigue originating from the central nervous system (5) and peripheral fatigue involving biochemical alterations within the muscles (6, 7). Moreover, classically, resistance and endurance exercises are thought to depend on type II and type I muscle fibers, respectively. Extensive research has elucidated the molecular and metabolic changes associated with these exercise types in a fiber-specific manner (8, 9). Given this, muscles like the gastrocnemius, which contain a mix of fiber types, have also been shown to be engaged in high-intensity exercise regimens and exhibit significant metabolic changes, particularly in key determinants of energetics such as NADH, ATP, P-Creatinine, and 5’ AMP-activated protein kinase (AMPK) (10, 11).

More importantly, recent papers have profiled the distinct metabolic response of skeletal muscle in response to exercise performed at different times of the day (2, 3, 12, 13). However, skeletal muscle intrinsic and extrinsic factors which are causal to such distinct metabolic adaptations and can potentially drive time of the day-dependent changes in the exercise capacity of the organism are still largely unanswered. Towards this, recent studies show that exercise performance in mice is circadian clock-regulated, as clock-disrupted genetic models (e.g., BAML1 and PER mutants) lack time-of-day performance variations (3). Approximately 4% of the skeletal muscle transcriptome in mice exhibits circadian rhythms, with genes involved in substrate utilization, storage, and muscle contraction (14, 15).

Mitochondria are crucial due to their role in circadian metabolism (16) and exercise capacity (17). Mitochondrial functions correlate strongly with exercise capacity, influencing energy demands, metabolic adaptation, calcium homeostasis, toxic intermediate clearance, and signaling pathway activation (18). Genetic and pharmacological perturbations targeting mitochondrial functions are linked to exercise capacity changes (19). Recent reports have also indicated that mitochondrial functions are regulated by core clock proteins including CLOCK and BMAL1 in skeletal muscle (20), though their impact on time-of-day-dependent exercise capacity remains unclear.

We hypothesize that mitochondrial function in skeletal muscle changes with the time of day, leading to differential metabolic and energetic adaptations in response to exercise performed at different times. In this study, our findings provide novel insights into the time-of-day-dependent regulation of exercise performance and underscore the importance of mitochondrial function in these processes.

## RESULTS

### Exercise capacity displays a time of the day dependent changes in the mouse

Towards unraveling the relationship between time of the day exercise capacity, metabolism, and mitochondrial energetics, wild type 2-4-month-old male mice were made to run on a treadmill as indicated **(Figure 1A)**. Even though several exercise paradigms have been employed in the literature (21), we chose a short-duration high-intensity exercise to minimize Zeitgeber (ZT)-dependent variation (which could otherwise be confounded by long-duration exercise). Mice were habituated to the treadmill for three days to negate the confounding effects of handling and stress associated with the novel arena. We found time of the day-dependent change in exercise capacity, which was assessed using total run length and time to fatigue, drastically decreased at ZT3 compared to ZT15 **(Figure S1A and 1B-C)**. This is consistent with many studies that have reported late active phase fatigue resistance including in humans (2–4). Given this, we proceeded with characterizing metabolic and mitochondrial underpinnings of exercise capacity at ZT3 and ZT15. Moreover, it is important to note that, contrary to the common assumption, findings in the literature point to the involvement of mixed muscle groups and outputs associated with mitochondrial functions in response to high-intensity exercise (9, 22).

**Figure 1:**
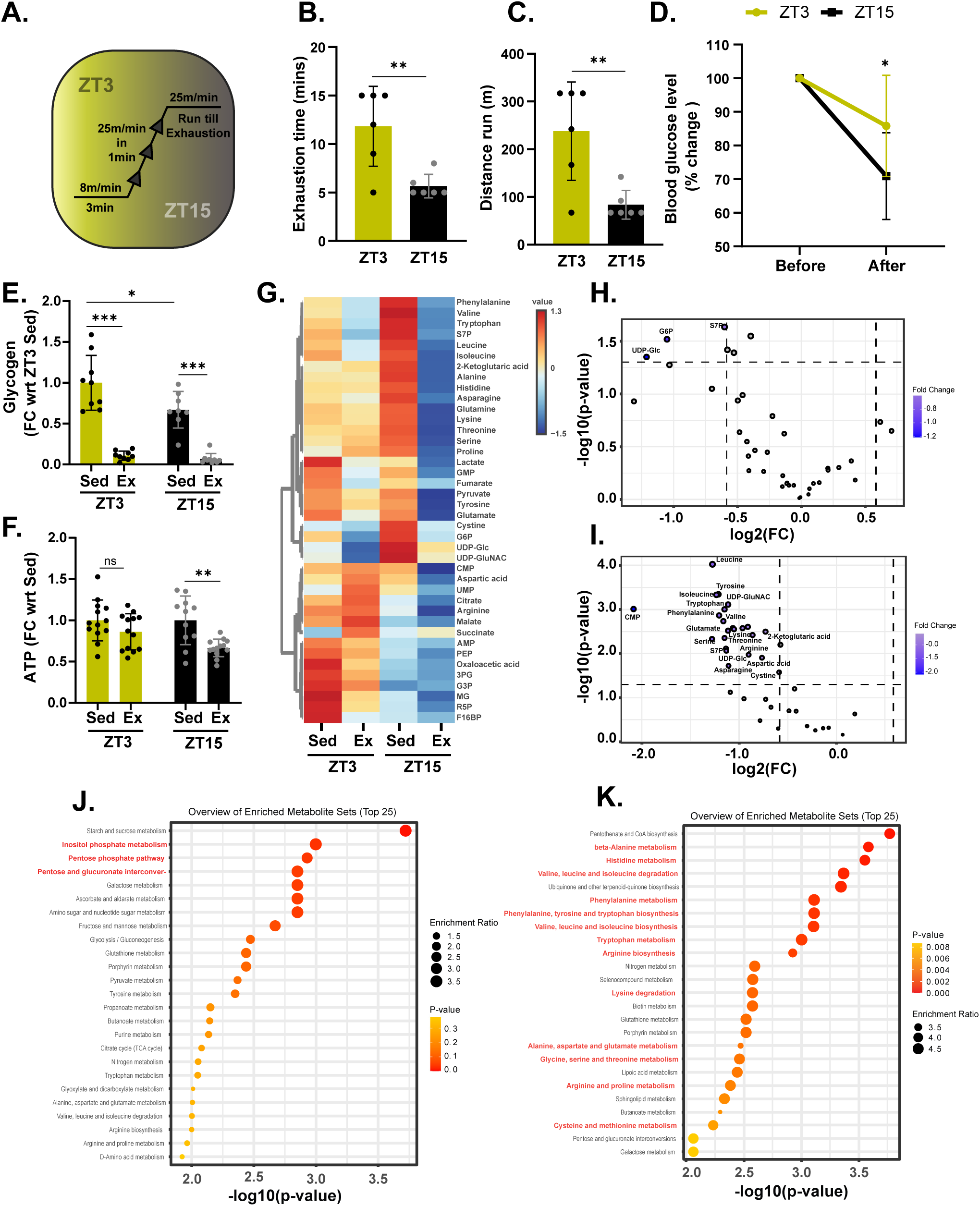
The metabolic and energetic profile of time of the day-dependent exercise is distinct in mice: (A) Schematic representation of the single-bout treadmill exercise paradigm. (B-C) Assessment of exercise-induced exhaustion: (B) total run time and (C) total run distance until exhaustion. Data are presented as bar graphs (n=6). (D) Blood glucose levels measured before and after exercise, with data plotted as the percentage reduction in blood glucose compared to pre-exercise levels (n=12-13). (E) Gastrocnemius glycogen levels in sedentary and post-acute exercise conditions at ZT3 and ZT15. The fold change relative to ZT3 sedentary levels is plotted as a bar graph (n=8-9). (F) ATP level measurements in gastrocnemius muscle isolated from sedentary and post-acute exercise animals at ZT3 and ZT15. The fold change relative to the corresponding sedentary animals at each time point is plotted as a bar graph (n=12-13). (G) Heatmap representation of metabolite levels in sedentary and exercised animals at ZT3 and ZT15. The scale represents the normalized concentration of metabolites (n=3). (H-I) Volcano plots depicting significantly altered metabolites post-exercise compared to corresponding sedentary controls at (H) ZT3 and (I) ZT15. The dotted horizontal and vertical lines indicate significance thresholds (p ˂ 0.05 and fold change ˃ 1.5, respectively). Names of significant metabolites are labeled (n=3). (J-K) KEGG pathway enrichment analysis of metabolites altered by exercise at (J) ZT3 and (K) ZT15. Significantly enriched pathways are highlighted in red. The enrichment score and p-value are indicated by size and color scales, respectively (n=3). Data in panels B-F are plotted as mean ± SD. Student’s *t*-test (B-D) and ANOVA (E-F) were used; \**p* ˂ 0.05, \*\**p* ˂ 0.01, \*\*\**p* ˂ 0.001, nonsignificant (ns) *p* ˃ 0.05.

Assaying for the blood glucose before and after exercise **(Figure 1D)** and measuring the respiratory exchange ratios (RER) **(Figure S1B, S1C, and S1D)** showed ZT-dependent change *vis-a-vis* substrate oxidation, as indicated. Additionally, gastrocnemius glycogen content was slightly lower at ZT15, and post-exercise, the muscle glycogen levels significantly decreased at both ZT3 and ZT15 **(Figure 1E).** Since these posited ZT-dependent and exercise-mediated metabolic plasticity, we performed muscle metabolomics as detailed in the methods section. As described in the introduction, gastrocnemius has been shown to respond to high-intensity exercise and display changes in key metabolites like NADH and phospho-creatinine (9, 10). This prompted us to unravel the time of the day-dependent metabolic and energetic rewiring in the mixed muscle group which was hitherto unknown.

It was interesting to note that the baseline muscle metabolome profiles at ZT3 and ZT15 were distinct with glycolytic intermediates and nucleotide metabolism prominently different in the sedentary conditions, in these two-time points **(Figure S1E and 1G)**. Following exercise, there was a significant change in the metabolome and the effect of exercise on the amino acid metabolism stood out, especially at ZT15 **(Figure 1H-K and Figure S1G)**. Based on the skeletal muscle metabolome profile and because muscle activity is inherently energy-demanding, we checked the total ATP from muscles isolated at ZT3 and ZT15. It was intriguing to find that while at ZT3 there was not much change in total ATP before and after exercise, there was a significant decrease post-exercise at ZT15 **(Figure 1F)**. This was consistent with the hypothesis based on the literature and also suggested a potential interplay between mitochondrial metabolism and energetics in driving ZT-dependent exercise adaptations and capacity in mice.

### Skeletal muscle mitochondrial OCR displays a time-of-the-day effect

The results described above led us to hypothesize mitochondrial metabolism and energetics as key determinants of time of day-dependent exercise capacity. Moreover, although muscle activity and mitochondrial functions have been independently shown to display circadian dependence (20), whether mitochondrial energetics contributes to muscle capacity in a time-of-the-day-dependent manner has not been investigated, thus far.

Towards this, we first assayed for oxygen consumption rates (OCR), electron transport chain (ETC) functions, and substrate oxidation from mitochondria isolated from muscles at ZT3 and ZT15 **(Figure 2A)**. We found a significant difference in mitochondrial respiration with a robust decrease in OCR at ZT15 when compared to ZT3, which was independent of Complex-I and II substrates **(Figure 2B-C and S2A-B)**. Notably, there was a marked reduction in state 3 respiration **(Figure S2A-B)** which corroborated with reduced ATP production in mitochondria isolated from ZT15 **(Figure 2E)**. We also found a consistent decrease in state U respiration at ZT15 **(Figure S2A-B)**. We also observed lower ETC flux as shown in **Figures 2D and S2C**, indicating a reduced bioenergetic potential at ZT15. Together these results reveal time of the day dependent intrinsic differences in gastrocnemius muscle mitochondrial functions.

**Figure 2:**
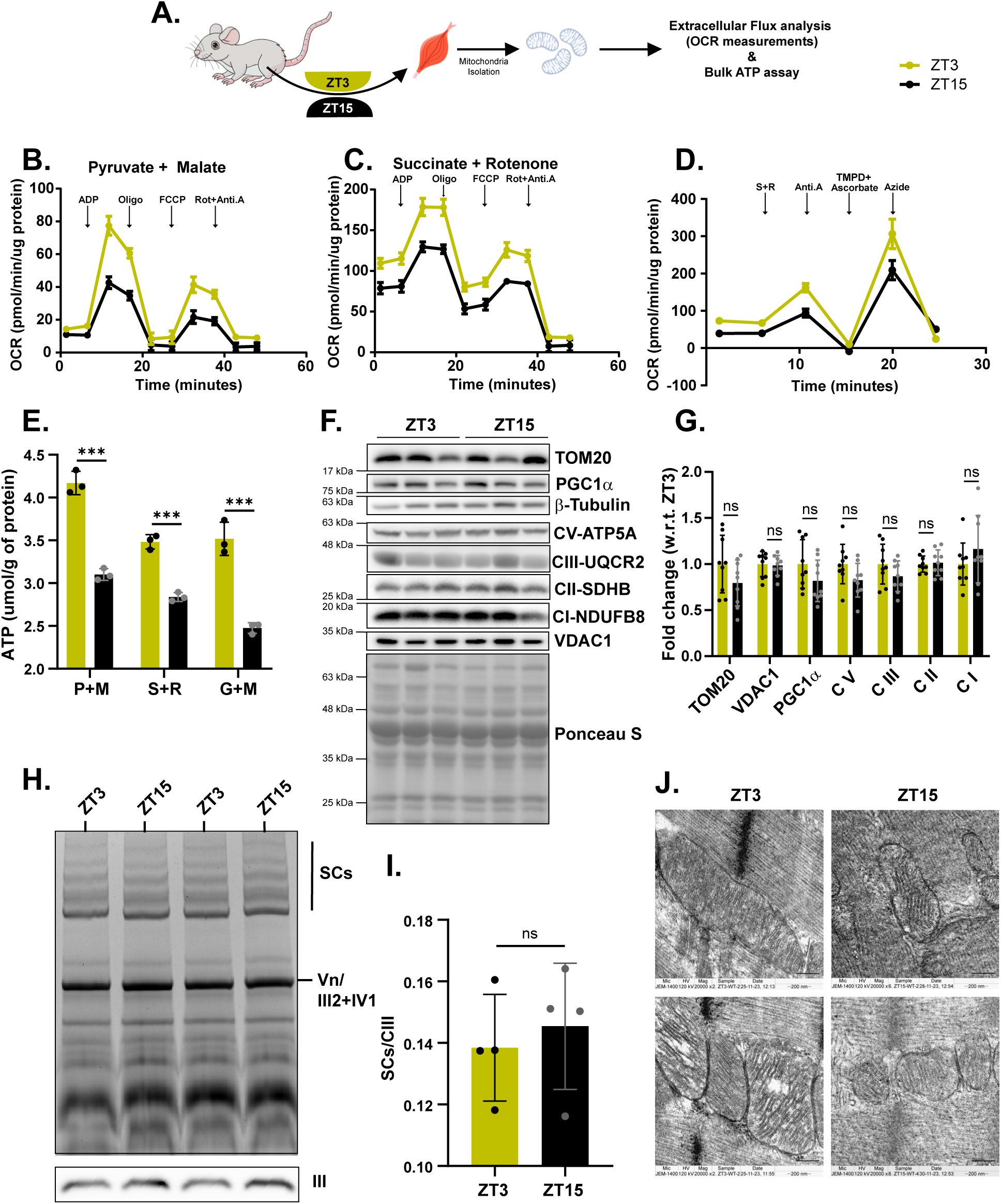
Skeletal muscle mitochondrial function changes in a time of the day-dependent manner: (A) Schematic representation of the oxygen consumption rate (OCR) measurement from isolated skeletal muscle mitochondria using the Agilent XFe24 Analyzer. (B-C) OCR time-course graphs for state respiration assays using isolated mitochondria from gastrocnemius muscle with substrates for (B) Complex I (Pyruvate + Malate) and (C) Complex II (Succinate + Rotenone) (N=3, n=5). The vertical arrows indicate the time points of compound injections: Oligo (Oligomycin), Rot+Anti.A (Rotenone + Antimycin A). (D) OCR time-course graph for the electron flow assay using isolated mitochondria from gastrocnemius muscle (N=3, n=10). The vertical arrow indicate the time points of compound injections: S+R (Succinate + Rotenone), Anti.A (Antimycin A), Azide (Potassium azide). (E) Quantification of ATP production. Data are represented as normalized ATP production in the presence of various substrates (N=3, n=3): P+M (Pyruvate + Malate), S+R (Succinate + Rotenone), G+M (Glutamate + Malate). (F-G) SDS-PAGE analysis of gastrocnemius muscle protein lysates collected at ZT3 and ZT15: (F) Representative immunoblots for mitochondrial markers (TOM20, VDAC1, PGC1α) and electron transport chain (ETC) complexes, with total protein stained with Ponceau S as a loading control (N=3, n=3). (G) Quantification of the immunoblots shown in (F), with data plotted as fold change relative to ZT3 (N=3, n=3). (H-I) Blue native gel electrophoresis analysis of mitochondrial supercomplexes in gastrocnemius muscle harvested at ZT3 and ZT15. (H) Representative images showing supercomplexes (SCs), Complex III (CIII), Complex IV (CIV), and Complex V (CV). Immunoblot for Complex III was used as a loading control (N=2, n=2). (I) Quantification of supercomplexes from the blue native gel images shown in (H), normalized to the Complex III immunoblot (N=2, n=2). (J) Transmission electron microscopy (TEM) analysis of gastrocnemius muscle harvested from mice at ZT3 and ZT15. Representative images are shown at 20,000x magnification with a scale bar of 200 nm. The data in panels B-E, F, and I are plotted as mean ± SD, and Student’s *t*-test was used to estimate statistical significance; \**p* ˂ 0.05, \*\**p* ˂ 0.01, \*\*\**p* ˂ 0.001, nonsignificant (ns) *p* ˃ 0.05.

### Mitochondrial functional differences are associated with mitochondrial morphology changes

While physiological context-dependent mitochondrial functions have been largely attributed to changes in mitochondrial content (23, 24), we did not find any alteration in markers that indicate mitochondrial biogenesis **(Figure 2F-G)**. This further pointed to mechanisms intrinsic to mitochondria that could dictate ZT-dependent mitochondrial functions. Notably, emerging literature has clearly shown that the changes in the ETC components, electron flow, and morphology are sufficient to affect mitochondrial output irrespective of biogenic potential (25–27). On assaying for the abundance of ETC components and supercomplexes, we were surprised to find no difference between isolated mitochondria from gastrocnemius muscle at ZT3 and ZT15 **(Figure 2F-2I)**.

Owing to the fact that mitochondrial morphology has surfaced as one of the major determinants of mitochondrial functions (28), we analyzed transmission electron micrographs of mitochondria from sections of gastrocnemius isolated at ZT3 and ZT15 **(Figure 2J)**. Computing mitometric parameters clearly indicated a change in the mitochondrial length, width, area, and aspect ratio **(Figure S2D-G)**, which is accompanied by a significant difference in the expression of *Mitofusin1 (Mfn1)* **(Figure S2H)**, a well-known regulator of mitochondrial fusion (29). In the absence of an overt change in biogenesis and/or ETC composition, we surmised that the subtle morphological difference coupled with mitochondrial metabolism may possibly be causal for the mitochondrial functional phenotype that we have observed, although this needs further validation.

### Sirt4^−/−^ mouse shows abrogated mitochondrial function

Differential substrate utilization and altered TCA flux are sufficient to drive mitochondrial functions, especially with regard to respiration and energetics (30, 31). Hence, based on the results presented above, we wanted to test if the time of the day exercise capacity is causally linked with mitochondrial functions. Towards this, we employed Sirtuin4 loss of function mice **(Figure S3A and B)**, as others and we have established its pivotal role as an upstream regulator of mitochondrial metabolism and energetics (31–34).

Histological analysis of skeletal muscles from wild-type (WT) and Sirtuin4 KO (Sirt4^−/−^) mice did not reveal any differences *vis-a-vis* cross-sectional area (CSA) and fiber types as indicated in **Figure 3A-3C and S3C**. Further, besides having similar muscle weights **(Figure S3E)**, molecular analysis for markers of trophism **(Figure 3D-E and S3F)** and fiber types **(Figure S3G)** not only ruled out gross abnormalities but also correlated well with the histological analysis. The absence of Sirt4 did not affect muscle glycogen content **(Figure S3H)** and glycogen metabolism genes **(Figure S3I)** at ZT3. However, it was interesting to note that mitochondria isolated from Sirt4^−/−^ muscles displayed lower OCR (state 3 and state U respirations), for both Pyruvate-Malate **(Figure 3F-G)** and Succinate **(Figure 3H-I)**, at ZT3. Notably, this was not associated with muscle circadian rhythm as indicated by the expression of clock genes at ZT3 and ZT15, which were comparable between WT and Sirt4^−/−^ mice **(Figure S3J)**.

**Figure 3:**
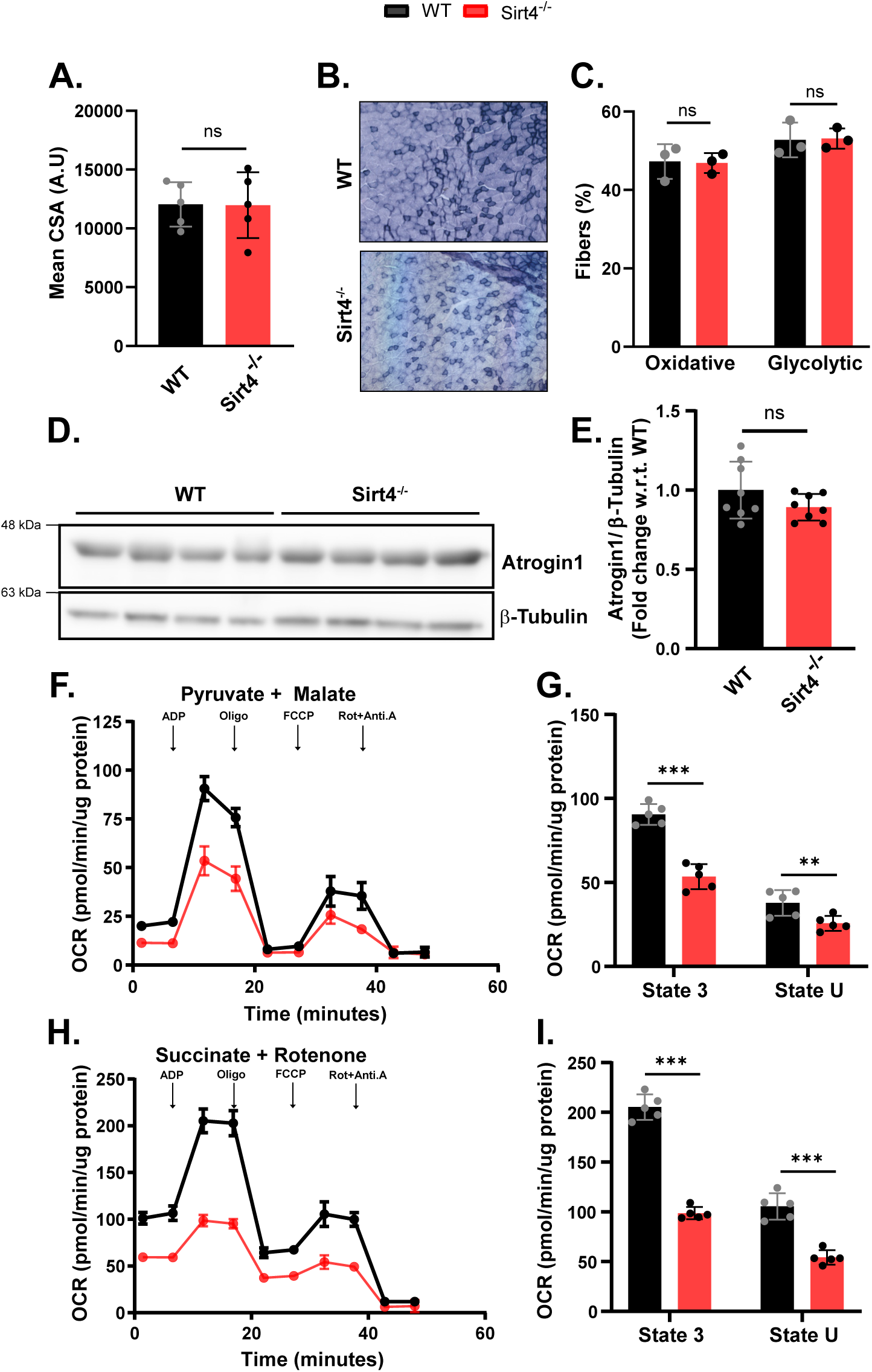
Loss of Sirt4 affects skeletal muscle mitochondrial functions without affecting trophism and fiber-type: (A) Quantification of the mean cross-sectional area of gastrocnemius muscle sections from wild-type (WT) and Sirt4^−/−^ mice, as shown in Figure S3C (N=2, n=5). (B-C) Fiber type profiling: (B) Representative images of succinate dehydrogenase (SDH) staining in WT and Sirt4^−/−^ muscle. (C) Bar graph representing the quantitative estimation of fiber type percentages based on SDH staining in WT and Sirt4^−/−^ muscle (N=2, n=3). (D-E) SDS-PAGE analysis of gastrocnemius muscle protein lysates from WT and Sirt4^−/−^ mice: (D) Representative western blot analysis for Atrogin-1 protein levels, normalized to β-tubulin. (E) Bar graph representing the quantification of normalized Atrogin-1 protein levels in WT and Sirt4^−/−^ muscle lysates (N=2, n=4). (F-I) OCR measurements from isolated mitochondria of the gastrocnemius muscle in WT and Sirt4^−/−^ mice: (F) OCR time-course graph using Complex I substrates (Pyruvate + Malate), and (H) OCR time-course graph using Complex II substrates (Succinate + Rotenone). (G) Quantification of State 3 and State U respiration for Pyruvate + Malate and (I) Succinate + Rotenone, based on the OCR time-course data. Vertical arrows indicate the injection points of the corresponding compounds: Oligo (Oligomycin), Rot + Anti.A (Rotenone + Antimycin A) (N=2, n=5). All data are plotted as mean ± SD, and Student’s *t*-test was used to estimate statistical significance; \**p* ˂ 0.05, \*\**p* ˂ 0.01, \*\*\**p* ˂ 0.001, nonsignificant (ns) *p* ˃ 0.05.

These results prompted us to investigate if mitochondria were affected as a function of time of day in Sirt4^−/−^ mice. While WT mice displayed ZT-dependent OCR, as shown earlier in **Figure 2**, the loss of Sirt4 seemed to abrogate this **(Figure 4A-F)**. Specifically, it was interesting to find that there was no difference in mitochondrial respiration **(Figure 4A-F)** between ZT3 and ZT15 in the absence of Sirt4. Moreover, the diminished mitochondrial output at both ZT3 and ZT15 responses in Sirt4^−/−^ mice mimicked the mitochondrial functions of WT mice at ZT15. Furthermore, transmission electron microscopy (TEM) images of Sirt4^−/−^ skeletal muscle sections revealed a loss of time-of-day-dependent changes in mitochondrial morphology **(Figure 4G and S4A-D)**. Additionally, molecular analysis of *Mitofusin1 (mfn1)* also displayed reduced levels at ZT3 in Sirt4^−/−^ skeletal muscle, possibly resulting in more fragmented mitochondria at both ZT3 and ZT15 **(Figure S4E)**. Moreover, there was no significant difference in mitochondrial content due to the loss of Sirt4 at both time points **(Figure 4H and S4F)**. In summary, the findings so far suggested that diurnal changes in mitochondrial morphology in WT animals are associated with circadian variations in mitochondrial functions, which are abrogated in Sirt4^−/−^ skeletal muscle.

**Figure 4:**
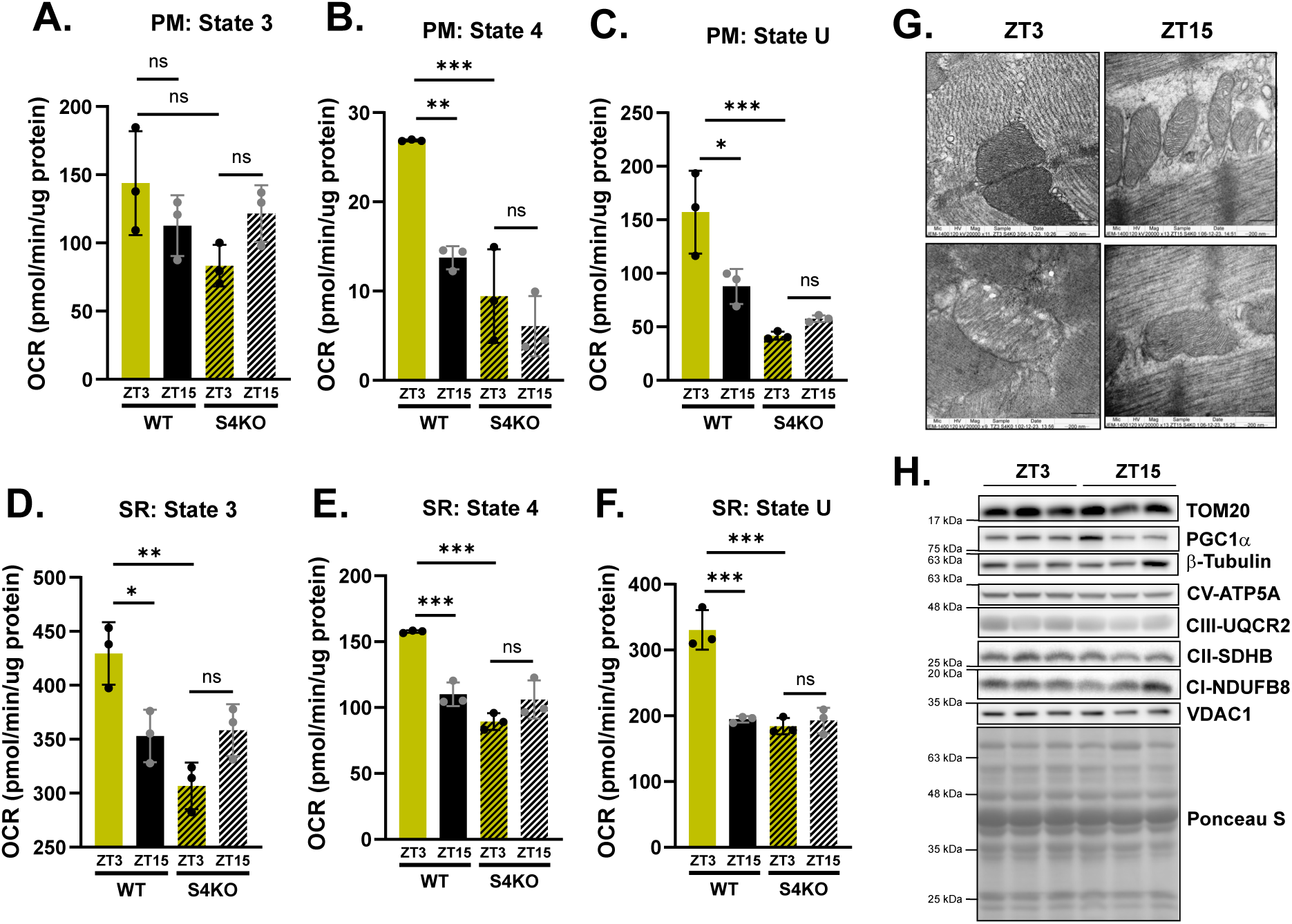
Loss of Sirt4 abrogates time of the day-dependent mitochondrial functions in skeletal muscle: (A-F) OCR measurements from isolated skeletal muscle mitochondria from wild-type (WT) and Sirt4^−/−^ mice at ZT3 and ZT15. Bar graphs represent OCR measurements using both (A-C) Complex I (Pyruvate + Malate) and (D-F) Complex II (Succinate + Rotenone) substrates: (A, D) State 3, (B, E) State 4, and (C, F) State U respiration were quantified for both substrates (N=3, n=3). (G) Transmission electron microscopy (TEM) analysis of gastrocnemius muscle from Sirt4^−/−^ mice at ZT3 and ZT15. Representative images are shown at 20,000x magnification with a scale bar of 200 nm (n=4). (H) SDS-PAGE analysis of gastrocnemius muscle protein lysates from Sirt4^−/−^ mice at ZT3 and ZT15. Representative immunoblots for mitochondrial markers (TOM20, VDAC1, PGC1α) and ETC complexes, with total protein stained with Ponceau S as a loading control (N=3, n=3). Data in panels A-F are plotted as mean ± SD, and two-way ANOVA (J) was used to estimate statistical significance; \**p* ˂ 0.05, \*\**p* ˂ 0.01, \*\*\**p* ˂ 0.001, nonsignificant (ns) *p* ˃ 0.05.

### Sirt4 Deficiency Abolishes Time-of-Day Effect on Exercise Capacity in Mice

Based on the results presented above, which clearly illustrated the importance of Sirt4 in determining time of the day mitochondrial functions, it was obvious for us to use Sirt4^−/−^ mice to ascertain mitochondrial causation *vis-a-vis* differential exercise capacity at ZT3 and ZT15. Toward this, we employed the same exercise paradigm as described in Figure 1A, and compared molecular and physiological parameters including exercise capacity between WT and Sirt4^−/−^ mice **(Figure 5A)**.

**Figure 5:**
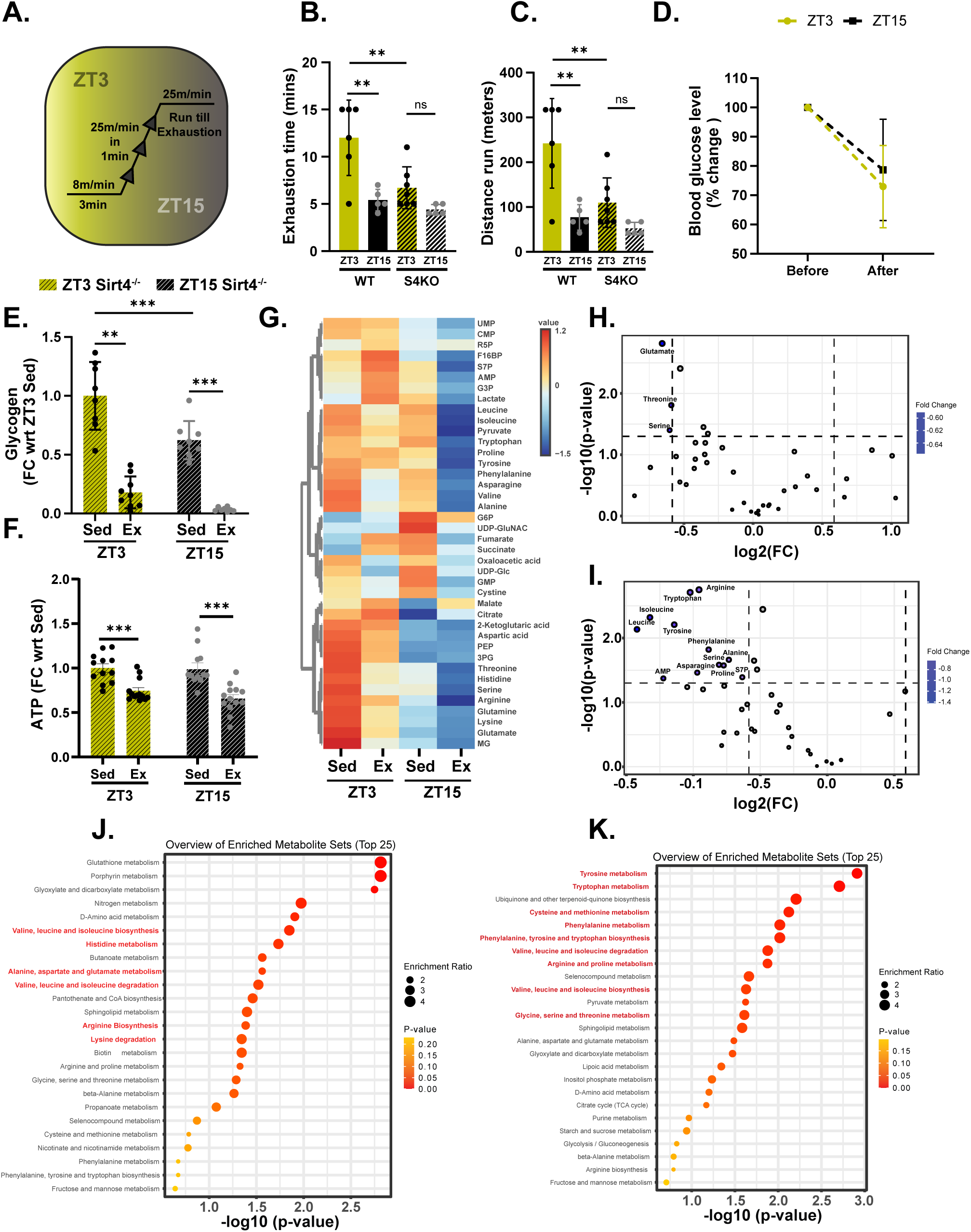
Diurnal differences in exercise capacity and associated metabolic and energetic adaptations are abolished in Sirt4^−/−^ mice: (A) Schematic representation of the single-bout treadmill exercise paradigm. (B-C) Assessment of exercise-induced exhaustion in WT and Sirt4^−/−^ mice by measuring (B) total run time and (C) total run distance (n=6). (D) Measurement of blood glucose levels before and after exercise. Data are plotted as the percentage reduction in blood glucose compared to pre-exercise levels at both time points (n=12-13). (E) Gastrocnemius glycogen levels in sedentary and post-acute exercise Sirt4^−/−^ animals at ZT3 and ZT15. The fold change relative to ZT3 sedentary levels is plotted (n=8-9). (F) Gastrocnemius ATP levels in sedentary and post-acute exercise Sirt4^−/−^ animals at ZT3 and ZT15. The fold change relative to the corresponding sedentary levels at each time point is plotted as a bar graph (n=12-13). (G) Heatmap representation of metabolite levels in sedentary and exercised Sirt4^−/−^ animals at ZT3 and ZT15. The scale represents normalized metabolite concentrations (n=3). (H-I) Volcano plot representation of metabolites significantly altered post-exercise compared to corresponding sedentary Sirt4^−/−^ animals at (H) ZT3 and (I) ZT15. The dotted horizontal and vertical lines indicate significance thresholds (p ˂ 0.05 and fold change ˃ 1.5, respectively). Names of significant metabolites are indicated (n=3). (J-K) KEGG pathway enrichment analysis of metabolites altered by exercise in Sirt4^−/−^ animals at (J) ZT3 and (K) ZT15. Significantly enriched pathways are highlighted in red. The enrichment score and p-value are indicated by size and color scales, respectively (n=3). Data in panels B-F are plotted as mean ± SD and the ANOVA test (E-F) was used to calculate statistical significance; \**p* ˂ 0.05, \*\**p* ˂ 0.01, \*\*\**p* ˂ 0.001, nonsignificant (ns) *p* ˃ 0.05.

We found a stark difference between these groups on both the measures of duration and total distance run until exhaustion **(Figure 5B-C)**. Notably, while WT mice displayed time of the day-dependent exercise capacity, loss of Sirt4 led to a drastic reduction in performance at both ZT3 and ZT15, which was comparable to WT ZT15 **(Figure 5B-C)**. Not surprisingly, this was associated with expected outcomes on circulating glucose **(Figure 5D)** and RER **(S5A-C)** unlike in the case of WT animals. However, exercise-induced glycogen utilization was slightly higher at ZT3 in Sirt4^−/−^ skeletal muscle than at ZT15 and interestingly mirrored what was observed in WT muscles **(Figure 5E)**.

Having ruled out trophic changes in the muscles and differential glycogen utilization as causal factors, we next checked for potential changes in ATP and metabolites. We found a significant difference between WT and Sirt4^−/−^ mice on assaying for muscle ATP at ZT3 but not at ZT15, which was consistent with our hypothesis **(Figure 5F)**. Further metabolome analysis on metabolites extracted from WT and Sirt4^−/−^ skeletal muscles at ZT3 and ZT15 also revealed distinct changes with respect to what was observed in the case of WT mice **(Figure 5G and S5D)**. It is important to note that exercise-mediated ZT-dependent change in muscle metabolome was abrogated in Sirt4^−/−^ mice **(Figure 5H-I)**. Specifically, Unlike in WT animals, amino acid metabolism seemed to be essential at both ZT3 and ZT15 post-exercise in Sirt4^−/−^ mice **(Figure 5H-K)**. Together, these results strongly support our hypothesis that time-of-day-dependent modulation in mitochondrial functions plays a crucial role in dictating the energetic and metabolic adaptations to exercise, consequently affecting peripheral exercise fatigue in gastrocnemius.

## DISCUSSION

Physical activity, particularly exercise, is fundamentally dependent on muscle and systemic metabolism, which are intrinsically linked to circadian rhythms. Evidence indicates that clock mechanisms within muscle tissue regulate its physiology, leading to a reciprocal relationship between exercise physiology and circadian rhythms in both rodents and humans (1). It is evident that athletic/sports performance exhibits time-of-day-dependent variations (35), although the underlying mechanisms, especially concerning exercise capacity and muscle energetics, remain poorly elucidated. In this regard, this study provides conclusive evidence that links muscle mitochondrial functions with time of the day-dependent exercise capacity, which was hitherto unknown. This is also relevant because it is important to ascertain if/how baseline muscle intrinsic ZT-dependent differences in metabolism and energetics are associated with exercise-induced fatigue.

Addressing circadian-dependent muscle functions and the long-term impact of time-of-day-dependent exercise is vital not only for fundamental scientific insights but also for its implications on human health. Although diverse exercise regimens and non-overlapping physiological measures have led to inconsistent interpretations, humans show improved short-duration maximal exercise performance in the afternoon under neutral climate conditions (4). Time-of-day dependent measurements across muscle groups including vastus lateralis and biceps femoris, during repeated sprints, have shown higher total work, peak power, and peak power decrement in the evening compared to the morning, even though this was not associated with any change in electromyography (EMG) activity (4, 36). Based on these findings, the field has posited that central factors may not significantly influence time-of-day-dependent variations vis-a-vis short-duration maximal exercise performance (4, 36).

Furthermore, exercise performance relies heavily on systemic metabolic adaptations, especially liver metabolic adaptations and liver-muscle crosstalk (12). Liver glycogenolysis and gluconeogenesis ensure continuous glucose supply to contracting muscles (37). Liver glycogen levels, which correlate with exercise duration, are higher at the late active phase as compared to the early active phase (3). Sato et al. (2022) showed that the skeletal muscle glycemic needs of mice exercising at ZT3 could be met by liver glycogen stores (12), this also aligns well with our blood glucose measurements and RER data indicating higher carbohydrate oxidation in the ZT3 exercise group (13). Conversely, mice exercising at ZT15, with smaller liver glycogen reserves, rely on alternative energy sources (12). This is supported by our metabolomics data showing higher amino acid catabolism in skeletal muscle at ZT15 and slightly lower RER values, suggesting reliance on proteins or fats for energy. Given these distinct systemic and muscle metabolic responses to exercise at different times of day, our study aimed to provide molecular insights into baseline time-of-day-dependent metabolic and mitochondrial factors that potentially influence muscle fatigue following a single bout of exercise.

Our results suggest that skeletal muscle mitochondria exhibit time-of-day-dependent changes in substrate oxidation and ATP production, potentially linked to mitochondrial morphology. Using Sirt4^−/−^ animals, we investigated the role of diurnal changes in mitochondrial function in regulating exercise capacity. The absence of Sirt4 significantly impaired mitochondrial functions without affecting other factors like muscle trophism, fiber type, glycogen metabolism, and the circadian system. This is important since core clock gene alterations affect both mitochondrial and non-mitochondrial factors (3, 38), making it difficult to dissect the specific role of circadian-dependent mitochondrial modulation in regulating muscle metabolism and exercise capacity. Furthermore, we observed that the loss of Sirt4 led to mitochondrial fragmentation at ZT3, resembling ZT15 WT mitochondria. The effect of Sirt4 on mitochondrial morphology is consistent with several studies performed in cell lines (39, 40). Consequently, Sirt4^−/−^ animals lost mitochondrial functional rhythmicity, resulting in a consistent decline in performance similar to ZT15 WT animals.

To assess the impact of lost mitochondrial rhythmicity on exercise performance, we compared exercise capacity between WT and Sirt4^−/−^ animals at ZT3 and ZT15. Our findings showed that Sirt4KO animal’s exercise capacity and metabolic/energetic adaptations no longer displayed time-of-day-dependent differences, resembling WT animals at ZT15. A targeted metabolomic approach revealed that amino acid metabolism is potentially regulated by mitochondrial functions in a time-of-day-dependent manner, influencing exercise capacity. Future investigations should explore how mitochondrial functions modulate exercise-induced amino acid metabolism and how amino acids contribute to exercise adaptations.

A significant limitation of our study is the focus on skeletal muscle. While it provides insights into the interplay between circadian rhythms, mitochondrial function, and exercise capacity, exercise performance is a systemic phenomenon influenced by various tissues and organs. Future research should adopt a comprehensive systemic approach to fully understand the dynamics governing circadian exercise physiology. Additionally, investigating the long-term impact of exercising at specific times of the day on mitochondrial functions, systemic metabolism, and the health benefits of exercise training will be an important area of study.

## MATERIALS AND METHODS

### Animals

The 129Sv mice were maintained under standard animal housing conditions, with a 12-hour light-dark cycle (lights on at 7 AM and off at 7 PM). Accordingly, 10 AM corresponds to ZT3 and 10 PM to ZT15. The housing temperature was maintained at 23°C. The animals were fed ad libitum with a standard chow diet. Littermates were housed together in a cage. WT (129Sv) and Sirt4^−/−^ (Sirt4tm1Fwa) mice were purchased from Jackson Laboratory. WT animals were crossed with Sirt4KO animals to generate Sirt4^+/−^ animals. These Sirt4^+/−^ animals were intercrossed for a minimum of 10 generations, after which Sirt4WT and Sirt4KO progeny from the Sirt4^+/−^ intercross were selected to establish Sirt4WT and Sirt4^−/−^ lines. Littermates from WT and KO breeding pairs were directly used for the experiments. The main breeding pairs for WT and Sirt4^−/−^ breeding were regularly replaced with new WT and KO animals derived from the Sirt4^+/−^ intercross. Male animals aged between 2 and 4 months were selected for the study. All procedures and the overall project were approved by the Institute Animal Ethics Committee (IAEC) and adhered to their guidelines. For ZT15 experiments, animals were handled and sacrificed under red light. Cervical dislocation was used for euthanasia. Following euthanization, the specified tissues were dissected and either snap-frozen in liquid nitrogen or processed for protein lysates and total RNA extraction.

### Treadmill exercise capacity

The exercise capacity of both WT and Sirt4^−/−^ mice was evaluated using metabolic treadmills (Columbus Instruments) at ZT3 (10 AM) and ZT15 (10 PM). Before the exercise capacity assessment, animals underwent a habituation period for three days, involving a 10-minute session at 8 m/min with a constant 5° incline on the treadmill. Mild electrical stimuli were applied to encourage mice to stay on the moving treadmill belt. For the exercise capacity measurement, animals performed a single bout of high-intensity exercise. The treadmill began at a speed of 8 m/min for 3 minutes, followed by an acceleration to 25 m/min in 1 minute. Subsequently, mice ran at a constant speed of 25 m/min until exhaustion criteria were met (receiving shocks ≥ 25/min). The time and distance until exhaustion were recorded to assess exercise capacity.

Additionally, blood glucose levels (measured with Accu-Check) were evaluated from tail blood both before placing the animals on the treadmill (pre-exercise) and post-exhaustion (post-exercise).

### Treadmill RER

To assess substrate oxidation during exercise, O_2_ consumption, and CO_2_ production were measured using indirect calorimetry in an Oxymax Comprehensive Lab Animal Monitoring System CLAMS open circuit system (Columbus Instruments, Columbus, OH, USA). WT and Sirt4^−/−^ mice underwent a 3-day habituation period on metabolic treadmills (Columbus Instruments) for 10 minutes at 8 m/min with a constant 5° incline. The mice received a mild electrical stimulus to encourage them to remain on the moving treadmill belt. O_2_ and CO_2_ gas sensors (Columbus Instruments) were calibrated before each test. Following the collection of resting gases, the treadmill was initiated at a speed of 8 m/min for 3 minutes, followed by an acceleration of 25 m/min in 1 minute. Subsequently, mice ran at a constant speed of 25 m/min for 10 minutes. After this period, the animals were allowed to recover for 15 minutes in the metabolic treadmills. The respiratory exchange ratio (RER) was calculated from the measured values of oxygen consumption (VO_2_) and carbon dioxide exhalation (VCO_2_). This ratio was used to estimate the relative contributions of fat and carbohydrate oxidation during exercise with the help of the following formulas (41):

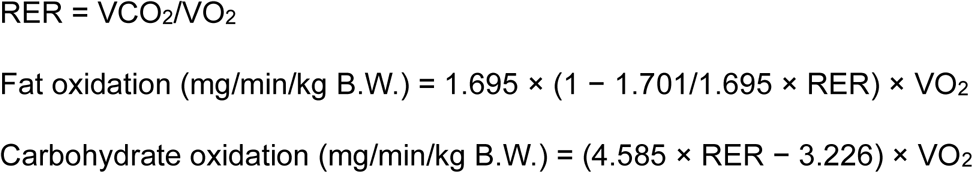

### Lysate preparation

Tissue lysates were generated using RIPA buffer (composition: 50mM Tris pH 8.0, 150mM NaCl, 0.1% SDS, 0.5% Sodium deoxycholate, 1% Triton X-100, 0.1% SDS, 1 mM PMSF, and Protease inhibitor cocktail). The lysates were centrifuged at 12,000 rpm (4°C/10 minutes) to pellet debris. Protein concentrations were quantified utilizing a BCA kit (Catalog number B9643) following the manufacturer’s instructions. Subsequently, the samples were subjected to boiling in an SDS gel loading buffer.

### Immunoblotting

Equal amounts of protein were run on an SDS-PAGE and transferred to polyvinylidene difluoride (PVDF) membranes. Ponceau S staining was performed, and a colorimetric image was captured using the GE Amersham Imager 600. After blocking with 5% milk, the membranes were probed with suitable primary and secondary antibodies, following standard protocols. Visualization of the bands was accomplished using a chemiluminescence detection kit (Catalog numbers 34580 and 34096) with the GE Amersham Imager 600. Ponceau S bands and Tubulin levels served as loading controls, as illustrated in the respective Figures 2-4. Band intensities were quantified utilizing ImageJ.

### Mitochondrial Supercomplexes assessment with blue native PAGE

100-150ug of isolated mitochondria was lysed in Solubilization Buffer (50mM NaCl, 50mM Imidazole-HCl, 2mM 6-AHA, 1mM EDTA) with 8g/g Digitonin (added from 10% digitonin solution in H_2_O) with a final concentration of 10ug/ul protein. The lysate was incubated in ice for 10min with mild vortexing pre and post-incubation. Lysate was spun at 20000xg, 20min, 4°C, and the supernatant was collected. 50% glycerol and 5% G-250 were added to a final concentration of 5% and 8g/g respectively. A 3-10% gradient gel was loaded in ice and run at 90V, 15mA, 30min for stacking after which Cathode Buffer A was replaced with Cathode Buffer B. The gel was further run at 300V, 15mA, 75min, post which the gel was incubated with Coomassie Blue G-250 stain on a slow rocker for 90min. Next, the gel was destained for 3-4 hours on a slow rocker or left undisturbed in destaining solution overnight, and imaged using a BioRad ChemiDoc Imaging System.

Buffers required:

Anode Buffer: 25 mM Imidazole.

Cathode Buffer: 50mM Tricine, 7.5mM Imidazole. Added 0.02% or 0.002% of 5% G-250 solution to make Cathode Buffers A and B respectively. Gel Buffer (3X): 75mM Imidazole, 1.5M 6-AHA.

Solutions:

5% Coomassie Blue G-250 in 500mM 6-AHA.

Acrylamide/Bis-Acrylamide Solution (49.5% T, 3% C).

### RNA extraction, RT-PCR, and qPCR analysis

Total RNA extraction from tissues was achieved using the Trizol reagent (Catalog number 15596018). Subsequently, 1 μg of RNA underwent reverse transcription using random hexamers (Catalog number N8080127) and a SuperScript-IV RT kit (Catalog number 18090200). Quantitative PCR (qPCR) was conducted employing KAPA SYBR® FAST Universal 2X qPCR Master Mix (Catalog number KK4602), and the Roche Light Cycler 480 II. RNA isolation, cDNA synthesis, and real-time PCR were performed following the manufacturer’s instructions. Levels of 18S and/or actin were utilized for normalization purposes.

### Tissue-ATP measurement

Gastrocnemius tissue was harvested and rapidly frozen in liquid nitrogen. ATP levels in the gastrocnemius tissue were quantified utilizing a luciferase-based assay, following the procedure outlined in (Ho et al., 2013). In brief, approximately 10mg of tissue was homogenized through crushing and heating in boiling water to facilitate ATP extraction. The ATP assay kit (Catalog number: FLLAM) was used to measure ATP levels in the extracted solution. The luminescence was measured according to the supplier’s protocol. Total ATP levels were estimated by referencing a standard curve and subsequently normalized to the total tissue protein.

### Tissue glycogen levels

Gastrocnemius tissue was collected and immediately frozen in liquid nitrogen. Gastrocnemius tissue samples weighing 10mg were homogenized on ice with 100μL of ice-cold sterile PBS buffer, followed by centrifugation at 12,000 rpm for 5 minutes to eliminate debris. For the assay, 10μl of the clear supernatant was utilized following the manufacturer’s protocol (Catalog number ab65620). Glycogen levels were normalized to the total tissue protein.

### Tissue sectioning and histological staining

The Gastrocnemius muscle was dissected and promptly snap-frozen in liquid nitrogen-cooled isopentane. Fresh-frozen sections (5-8 µm) were subsequently prepared on a Cryostat and stored at −80°C until needed. For histological examinations using Hematoxylin-Eosin (H&E)-staining, sections were dried at 37°C for 30-45 minutes. Afterward, routine H&E staining was performed on the sections, and the images were captured using an Olympus microscope at 10x magnification. The cross-sectional area was calculated in ImageJ and averaged for each mouse.

To determine fiber-type composition using Succinate Dehydrogenase (SDH) activity, sections were again dried at 37°C for 30-45 minutes. The dried sections were incubated with SDH incubating solution (1mg/ml nitroblue tetrazolium, 50mM Tris-buffer pH7.4, 0.5mM MgCl2, 4mg disodium succinate) for 1 hour at 37°C. The stained sections were washed with water for 5 minutes and air-dried. Subsequently, the sections were mounted using glycerine jelly and imaged with an Olympus microscope at 10x magnification. Fiber type was manually quantified in ImageJ, distinguishing between dark blue (oxidative fibers) and light blue (glycolytic fibers).

### Mitochondrial isolation

Animals were sacrificed at ZT3 or ZT15. Immediately post-sacrifice, the gastrocnemius muscle was excised, and all subsequent processing was carried out either on ice or at 4°C. All reagents and equipment were pre-chilled in ice. The muscle was minced in the Mitochondria Isolation Buffer (composition: 70 mM Sucrose, 220 mM Mannitol, 10 mM HEPES, 1 mM EGTA, 2mg/ml BSA, pH 7.2). Subsequently, a Teflon-Glass homogenizer (motorized) was utilized to homogenize the muscle with 15-18 strokes. The homogenate underwent two rounds of centrifugation at 800xg for 10 minutes to pellet nuclei and tissue debris. The resulting supernatant was further subjected to two rounds of centrifugation at 12,000xg for 10 minutes to obtain the mitochondrial pellet. This pellet was then resuspended in Mitochondrial Assay Solution (composition: 70mM Sucrose, 220mM Mannitol, 10mM KH2PO4, 2mM HEPES, 1mM EGTA, 5mM MgCl2, 4mg/ml BSA, pH 7.2) for all subsequent assays.

### Mitochondrial state respiration

The mitochondrial state respiration assay was conducted by measuring the Oxygen Consumption Rate (OCR) using a Seahorse Xfe24 Analyzer. After resuspending the pellet, protein estimation was performed, and 3-5ug of mitochondria were added to each well of a Seahorse plate, followed by centrifugation at 2000xg, 4°C for 20 minutes. Substrates were added into each well and the volume was made up to a total volume of 500ul (with MIB), and the plate was incubated in a non-CO_2_ incubator at 37°C for 8 minutes. For complex I-dependent respiration, Pyruvate/Malate (7.5mM/2.5mM) served as the substrate, while Succinate/Rotenone (10mM/2uM) was used for complex II-dependent respiration. Subsequently, sequential injections of ADP (100uM) (State III), Oligomycin (4uM) (State IV), FCCP (4uM) (State U), and Rotenone/Antimycin A (2uM/4uM) were administered to assess the corresponding state respiration.

### Mitochondrial ATP assay

The resuspended mitochondrial pellet was subjected to protein estimation (with BCA), and subsequently, 3-5ug of mitochondria were utilized for the assay. Mitochondria were incubated with complex-specific substrates and ADP in 1.5ml microcentrifuge tubes at 37°C for 30 minutes. After incubation, the tubes were centrifuged at 12,000xg, 4°C for 10 minutes to pellet the mitochondria. The resulting supernatant was used to measure ATP levels using a luciferase/luciferin-based assay system following the manufacturer’s protocol (Catalog number FLAAM).

### Electron Flow Assay (ETF assay)

The ETF assay was conducted following the protocol outlined by (Radogna et al., 2021) utilizing a Seahorse Xfe24 Analyzer. In brief, the resuspended pellet underwent protein estimation, and subsequently, 3-5ug of mitochondria were added to each well of a Seahorse plate and spun at 2000xg, 4°C for 20 minutes. Substrates were introduced into each well up to a total volume of 500ul, and the plate was incubated in a non-CO2 incubator at 37°C for 8 minutes. The basal media for the assay consisted of Pyruvate/Malate (7.5mM/2.5mM) (C-I-III-IV) with 4uM FCCP. This was followed by sequential injections of Succinate/Rotenone (10mM/2uM) (CII-III-IV), Antimycin A (4uM), TMPD/Ascorbate (100uM/10mM), and Potassium Azide (50mM) (CIV).

### Transmission Electron Microscopy (TEM)

The TEM experiment was performed with the help of the Electron Microscope facility, ACTREC, Tata Memorial Centre, Kharghar, Navi Mumbai.

The red part of the gastrocnemius muscle was cut into pinhead-sized pieces and fixed in 3% glutaraldehyde for 2 hours at 4°C, then post-fixed in 1% osmium tetroxide (OsO4) for 1 hour at 4°C. Care was taken to use the same red part of the muscle for fixation across samples. The tissues were block-stained in 2% uranyl acetate, followed by dehydration in an ascending alcohol series for 15 minutes each and finally with acetone. The tissues were infiltrated with Araldite resin (A and B), embedded in Araldite resin, and polymerized at 60°C for 48 hours. Ultrathin sections of 70 nm were cut using an ultramicrotome (Leica UC7), collected on 200 mesh copper grids, and contrasted with lead citrate. Images were acquired using a transmission electron microscope (JEM 1400 Plus, JEOL, Japan) at 120 kV. Image analysis was performed with the ImageJ application, quantifying around 200-250 mitochondria across four independent animals at both ZT3 and ZT15.

### Metabolomics

#### (a) Metabolites Extraction

Gastrocnemius muscle tissue (50 mg) from Sedentary or exercised animals was homogenized (3x, 20s) with a chilled tube on ice. One milliliter of cold extraction buffer (Ethanol: H2O, 4:1 v/v) was added. Metabolite extraction employed an internal control (13C4-aspartate) to normalize replicates, ensuring efficient extraction. Intracellular metabolites were extracted with 75% absolute ethanol, vortexed, and kept on ice. The process was repeated for 5 cycles, followed by incubation at −20°C for 1 hour and centrifugation at 16,000 g at 4°C for 15 minutes. The supernatant was transferred to a fresh tube, centrifuged again, and aliquots (300 µl) were dried using a speed vacuum. Samples were stored at −80°C until LC-MS/MS analysis. The peak intensity data was normalized with tissue weight and internal standard. The normalized data was further analyzed online on Metaboalnlyst 6.0.

#### (b) Metabolite profiling

Metabolites were detected and analyzed using highly quantitative LC-MS/MS, based on prior methods as described (Walvekar et al 2018 https://www.ncbi.nlm.nih.gov/pmc/articles/PMC6171562/). LC-MS/MS analysis utilized an AB SCIEX QTRAP 5500 with Waters Acquity UPLC system. All methods utilized a synergi 4-µm Fusion-RP 80 Å (150 × 4.6 mm) LC column. Targeted metabolite profiling, peak identification, and data processing were performed using Multi Quant software (version 3.0.1). LC-MS/MS parameters are provided in Table 1.

#### (c) Targeted Amino Acid Profiling

The amino acid analysis in positive polarity mode employed a synergi 4-µm Fusion-RP 80 Å (150 × 4.6 mm) LC column. Solvent A was water with 0.1% formic acid, and Solvent B was methanol with 0.1% formic acid.

#### (d) TCA Metabolites Profiling

Intracellular metabolites from gastrocnemius muscle tissue were extracted and dried. O-benzylhydroxylamine (OBHA) derivatization was performed for TCA cycle metabolites with functional carbonyl and carboxyl groups. The solvent system and gradients for separation are described. LC-MS/MS parameters are provided in Table1.

#### (e) Sugar Phosphate Metabolites Profiling

Intracellular metabolites from gastrocnemius muscle tissue were extracted and dried. Solvent systems and gradients for separation in negative polarity mode are provided. LC-MS/MS parameters are in Table 1. Details of the LC-MS/MS methods, along with metabolite information are available in Adhish W. et al., 2018 (42).

### Quantification and Statistical Analysis

The data are presented as means ± standard deviation (SD). Statistical analyses were conducted using Microsoft Excel (2016) and GraphPad Prism (version 6.0). Significance was determined through Student’s t-test and ANOVA. A p-value of ≤ 0.05 was considered statistically significant. Symbol notations for significance levels are as follows: *p ˂ 0.05; **p ˂ 0.01; ***p ˂ 0.001.

## Acknowledgments

We thank Dr. Kalidas Kohale and Dr. Shital Suryavanshi (TIFR-AH) and Dr. Suraj Ingle, Dr. Sagar Tarte, Ms. Ritika Gupta, Mr. Chetan Sable (National Facility for Gene Function in Health and Disease, IISER Pune) for help with the animal experiments. Dr. Gayathri Narayanappa and Dr. Srinivas Bharath (NIMHANS, Bangalore) for their help regarding skeletal muscle sectioning and imaging. ACTREC EM facility for their help with mitochondrial TEM image acquisition. inSTEM/NCBS/CCAMP mass spectrometry facility for instrument support for LC-MS/MS. We extend our acknowledgment to UK laboratory members for their critical input and discussions during the study.

## Funding Sources

This study was supported by the Department of Atomic Energy, Government of India, under Project Identification No. RTI4007 through TIFR under the grants numbers 19P0116 and 19P0911, Department of Science and Technology JCB/2022/000036, Department of Biotechnology (BT/PR29878/PFN/20/1431/2018) to U.K. DBT-Wellcome Trust India Alliance Senior Fellowship (IA/S/21/2/505922) to SL, S. Ramachandran National Bioscience Award for Career Development, Dept. of Biotechnology, to SL. We also thank the National Facility for Gene Function in Health and Disease (supported by a grant from the Department of Biotechnology, Government of India; BT/INF/22/ SP17358/2016) at IISER Pune for maintaining and providing mice used in this study.

## Supporting information

This article contains supporting information

## Data and materials availability

All data needed to evaluate the conclusions in the paper are present in the paper and/or the Supplementary Materials.

## Conflict of interest

The authors declare no conflict of interest.

---

**Figure S1.**
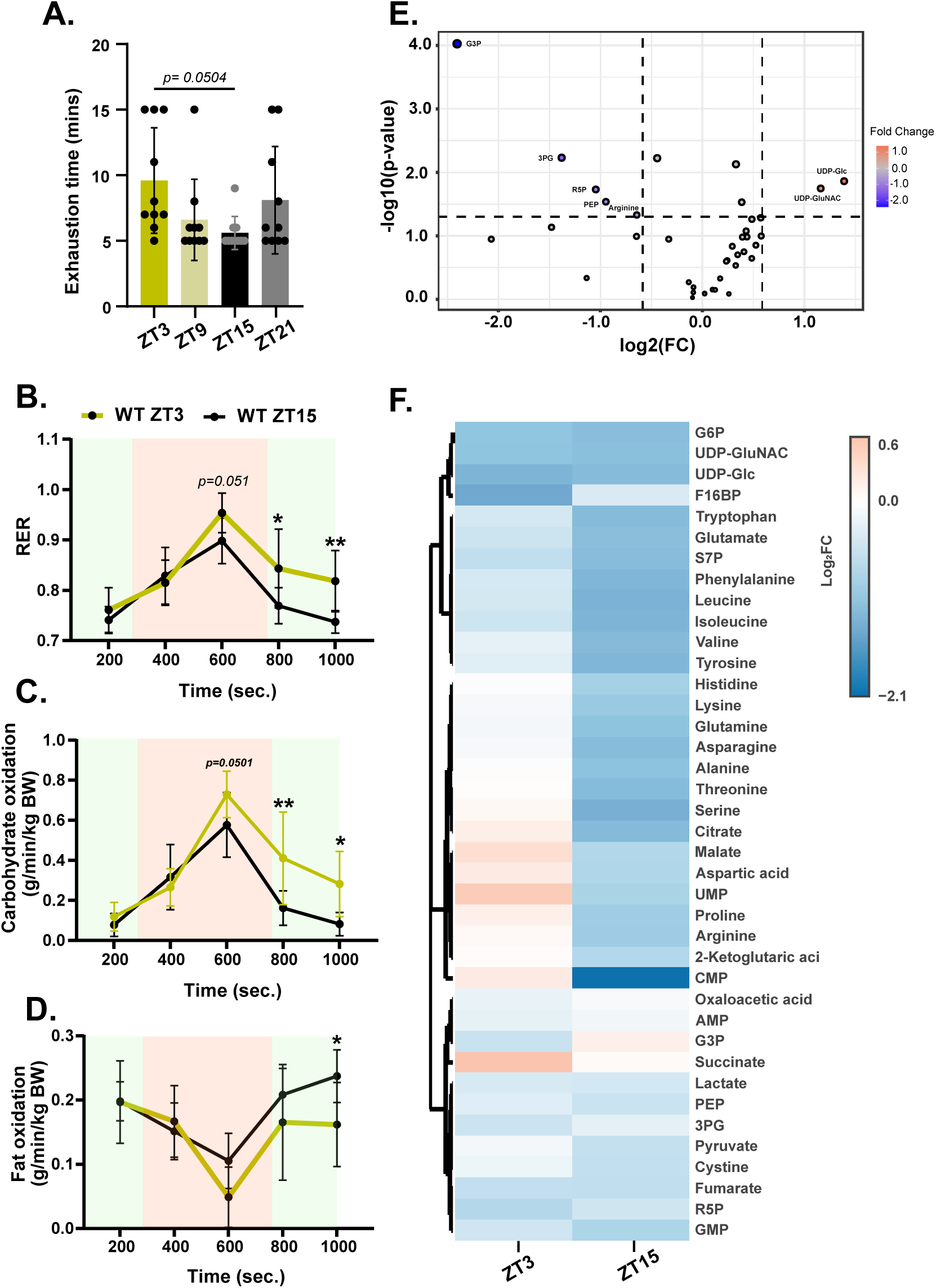

**Figure S2.**
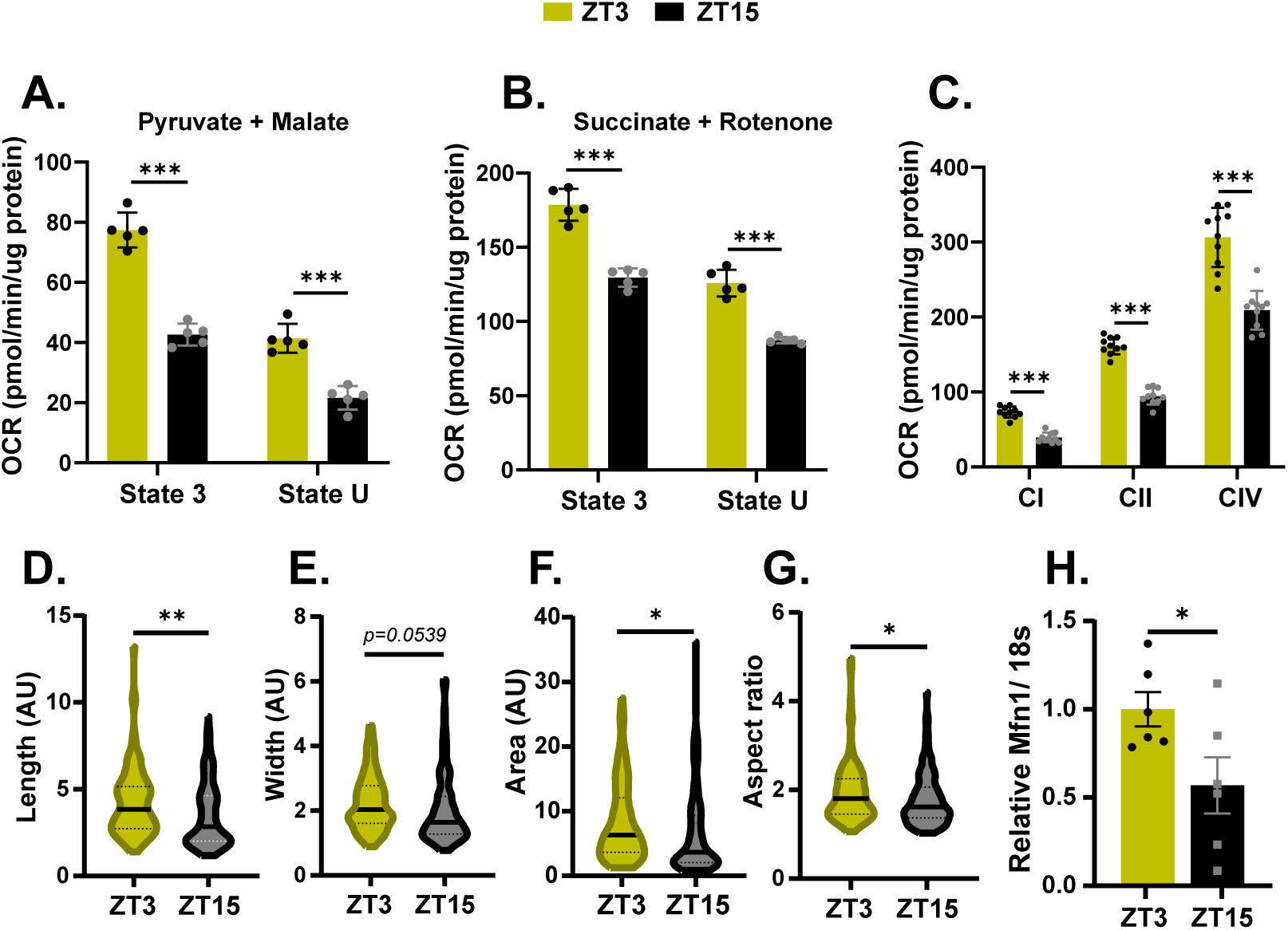

**Figure S3.**
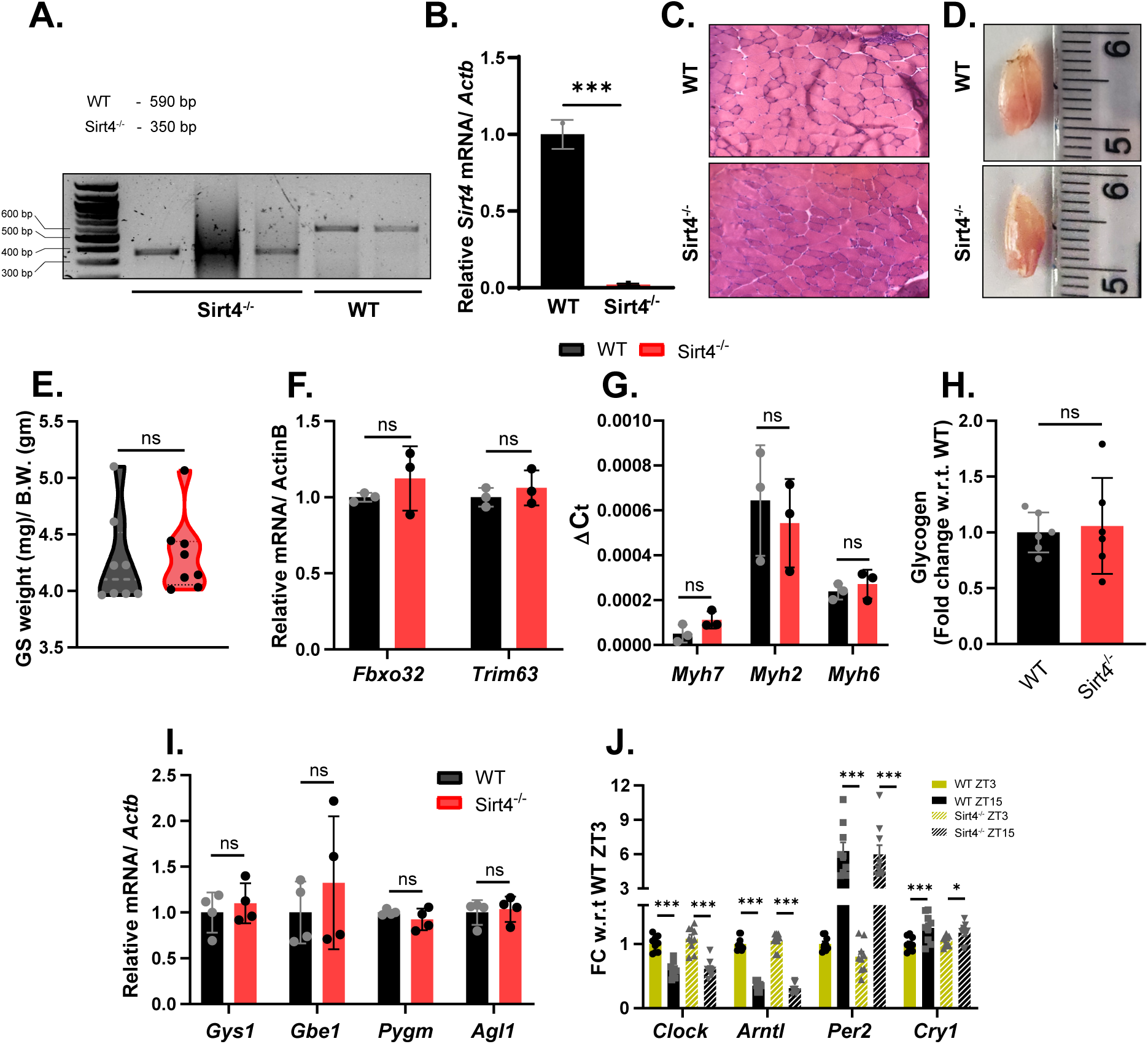

**Figure S4.**
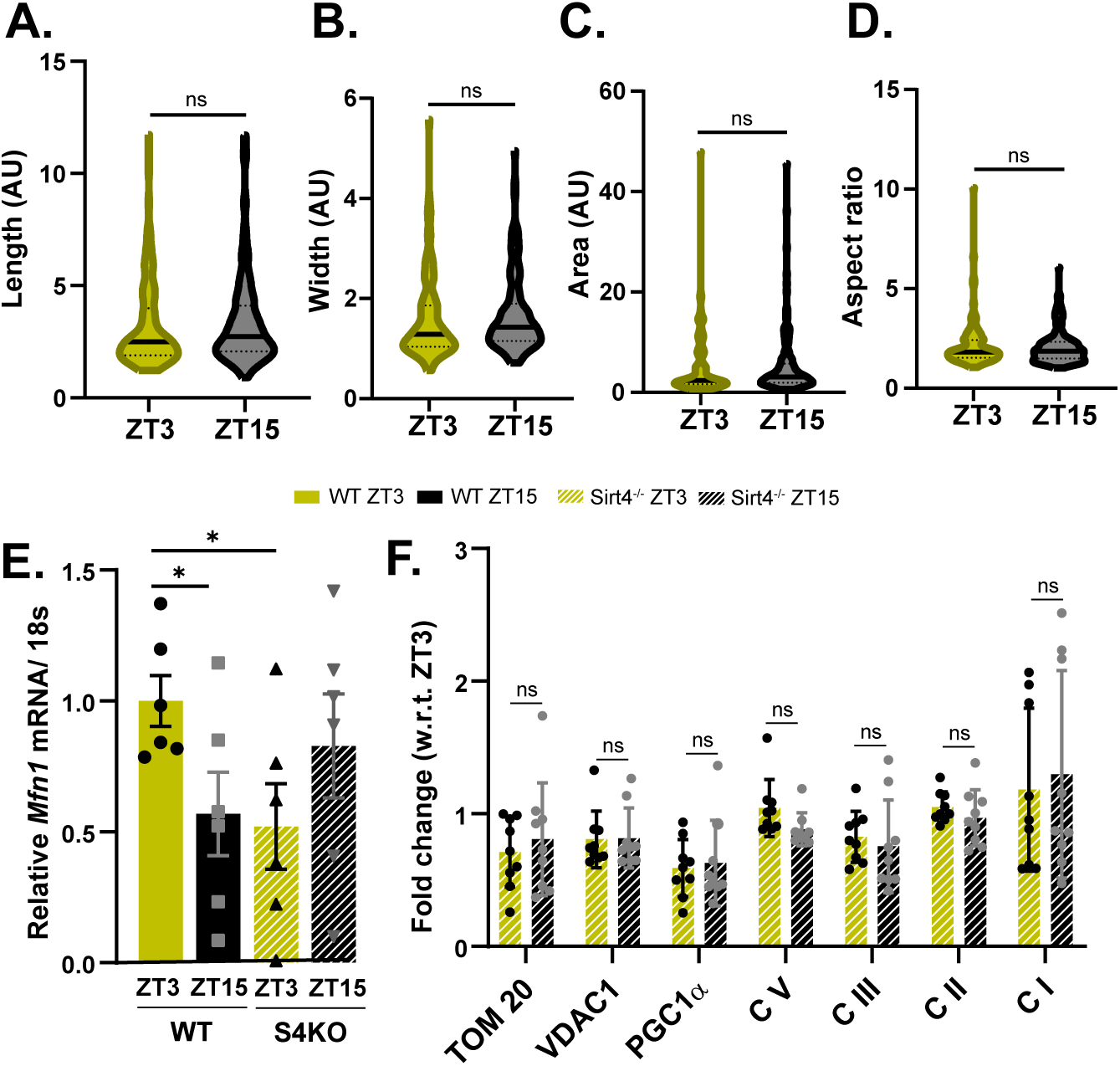

**Figure S5.**
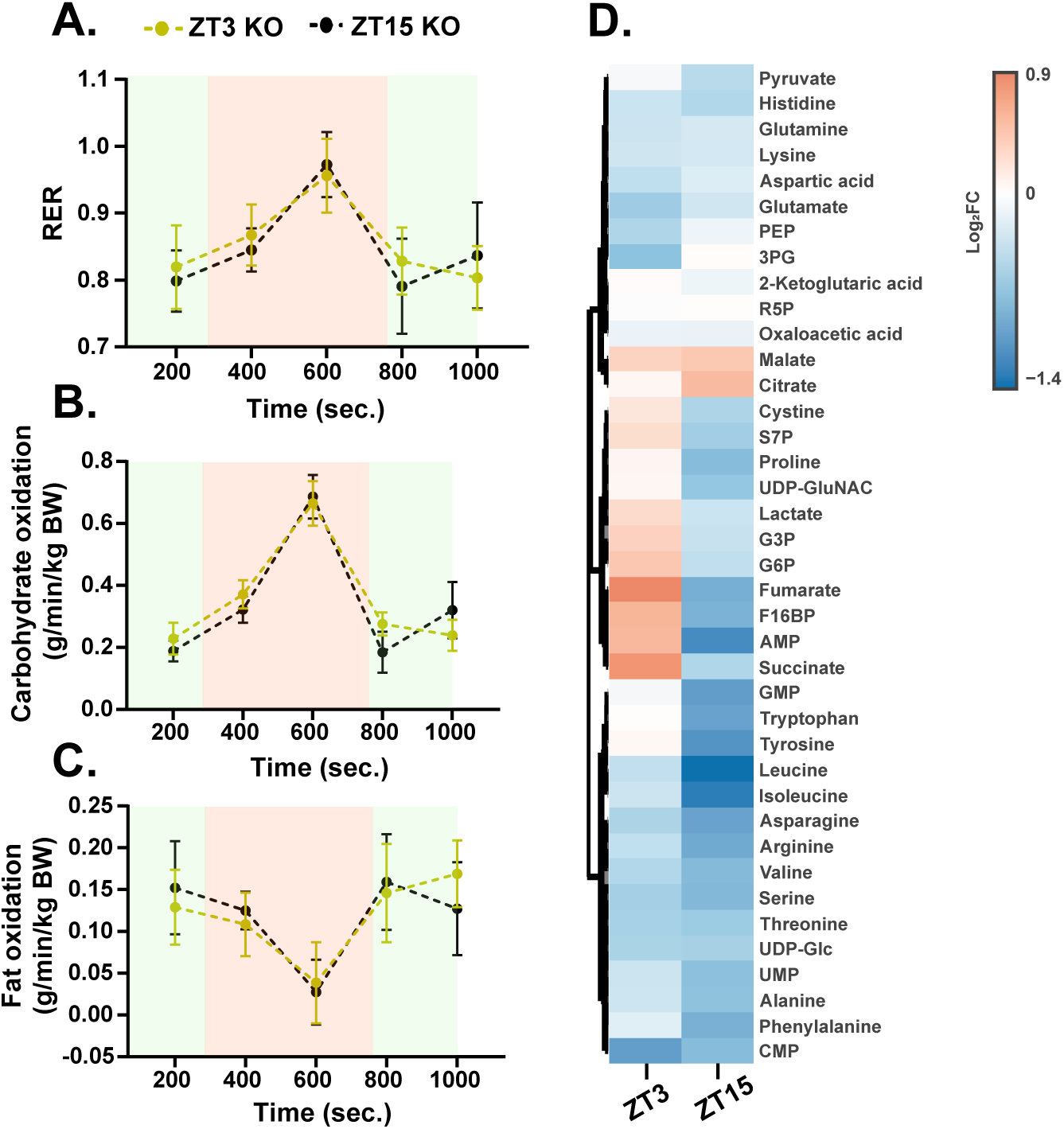

## Notes

### Competing Interest Statement

ARUMDA has received CSR funding from Hindustan Unilever Ltd. But the results presented in this study have no conflict of interest vis-a-vis this funding.

## REFERENCES

1. D. Duglan, K. A. Lamia, Clocking In, Working Out: Circadian Regulation of Exercise Physiology. Trends Endocrinol Metab 30, 347–356 (2019).

2. S. Ezagouri et al., Physiological and Molecular Dissection of Daily Variance in Exercise Capacity. Cell Metab 30, 78–91.e74 (2019).

3. Y. Adamovich et al., Clock proteins and training modify exercise capacity in a daytime-dependent manner. Proc Natl Acad Sci U S A 118 (2021).

4. G. G. Mirizio, R. S. M. Nunes, D. A. Vargas, C. Foster, E. Vieira, Time-of-Day Effects on Short-Duration Maximal Exercise Performance. Sci Rep 10, 9485 (2020).

5. Y. Yang et al., Exercise-Induced Central Fatigue: Biomarkers, and Non-Medicinal Interventions. Aging Dis (2024).

6. H. J. Green, Mechanisms of muscle fatigue in intense exercise. J Sports Sci 15, 247–256 (1997).

7. J. F. Tornero-Aguilera, J. Jimenez-Morcillo, A. Rubio-Zarapuz, V. J. Clemente-Suárez, Central and Peripheral Fatigue in Physical Exercise Explained: A Narrative Review. Int J Environ Res Public Health 19 (2022).

8. P. Krustrup, K. Söderlund, M. Mohr, J. González-Alonso, J. Bangsbo, Recruitment of fibre types and quadriceps muscle portions during repeated, intense knee-extensor exercise in humans. Pflugers Arch 449, 56–65 (2004).

9. J. M. Ren, J. Henriksson, A. Katz, K. Sahlin, NADH content in type I and type II human muscle fibres after dynamic exercise. Biochem J 251, 183–187 (1988).

10. A. Ratkevicius, M. Mizuno, E. Povilonis, B. Quistorff, Energy metabolism of the gastrocnemius and soleus muscles during isometric voluntary and electrically induced contractions in man. J Physiol 507 (Pt 2), 593–602 (1998).

11. T. E. Jensen et al., EMG-normalised kinase activation during exercise is higher in human gastrocnemius compared to soleus muscle. PLoS One 7, e31054 (2012).

12. S. Sato et al., Atlas of exercise metabolism reveals time-dependent signatures of metabolic homeostasis. Cell Metab 34, 329–345.e328 (2022).

13. S. Sato et al., Time of Exercise Specifies the Impact on Muscle Metabolic Pathways and Systemic Energy Homeostasis. Cell Metab 30, 92–110.e114 (2019).

14. R. Zhang, N. F. Lahens, H. I. Ballance, M. E. Hughes, J. B. Hogenesch, A circadian gene expression atlas in mammals: implications for biology and medicine. Proc Natl Acad Sci U S A 111, 16219–16224 (2014).

15. J. J. McCarthy et al., Identification of the circadian transcriptome in adult mouse skeletal muscle. Physiol Genomics 31, 86–95 (2007).

16. G. Manella, G. Asher, The Circadian Nature of Mitochondrial Biology. Front Endocrinol (Lausanne) 7, 162 (2016).

17. K. E. Conley, Mitochondria to motion: optimizing oxidative phosphorylation to improve exercise performance. J Exp Biol 219, 243–249 (2016).

18. A. S. Monzel, J. A. Enríquez, M. Picard, Multifaceted mitochondria: moving mitochondrial science beyond function and dysfunction. Nat Metab 5, 546–562 (2023).

19. P. M. Schaefer et al., Mitochondrial mutations alter endurance exercise response and determinants in mice. Proc Natl Acad Sci U S A 119, e2200549119 (2022).

20. J. L. Andrews et al., CLOCK and BMAL1 regulate MyoD and are necessary for maintenance of skeletal muscle phenotype and function. Proc Natl Acad Sci U S A 107, 19090–19095 (2010).

21. S. Marcaletti, C. Thomas, J. N. Feige, Exercise Performance Tests in Mice. Curr Protoc Mouse Biol 1, 141–154 (2011).

22. D. E. Kristensen et al., Human muscle fibre type-specific regulation of AMPK and downstream targets by exercise. J Physiol 593, 2053–2069 (2015).

23. A. Costanzini et al., Mitochondrial Mass Assessment in a Selected Cell Line under Different Metabolic Conditions. Cells 8 (2019).

24. A. Garnier et al., Coordinated changes in mitochondrial function and biogenesis in healthy and diseased human skeletal muscle. Faseb j 19, 43–52 (2005).

25. I. R. Lanza et al., Chronic caloric restriction preserves mitochondrial function in senescence without increasing mitochondrial biogenesis. Cell Metab 16, 777–788 (2012).

26. D. van Moorsel et al., Demonstration of a day-night rhythm in human skeletal muscle oxidative capacity. Mol Metab 5, 635–645 (2016).

27. N. Baker, J. Patel, M. Khacho, Linking mitochondrial dynamics, cristae remodeling and supercomplex formation: How mitochondrial structure can regulate bioenergetics. Mitochondrion 49, 259–268 (2019).

28. M. Liesa, O. S. Shirihai, Mitochondrial dynamics in the regulation of nutrient utilization and energy expenditure. Cell Metab 17, 491–506 (2013).

29. A. Santel et al., Mitofusin-1 protein is a generally expressed mediator of mitochondrial fusion in mammalian cells. J Cell Sci 116, 2763–2774 (2003).

30. N. M. Vacanti et al., Regulation of substrate utilization by the mitochondrial pyruvate carrier. Mol Cell 56, 425–435 (2014).

31. E. Shaw et al., Anabolic SIRT4 Exerts Retrograde Control over TORC1 Signaling by Glutamine Sparing in the Mitochondria. Mol Cell Biol 40 (2020).

32. L. Ho et al., SIRT4 regulates ATP homeostasis and mediates a retrograde signaling via AMPK. Aging (Albany NY) 5, 835–849 (2013).

33. N. Nasrin et al., SIRT4 regulates fatty acid oxidation and mitochondrial gene expression in liver and muscle cells. J Biol Chem 285, 31995–32002 (2010).

34. D. Tomaselli, C. Steegborn, A. Mai, D. Rotili, Sirt4: A Multifaceted Enzyme at the Crossroads of Mitochondrial Metabolism and Cancer. Front Oncol 10, 474 (2020).

35. R. Lok, G. Zerbini, M. C. M. Gordijn, D. G. M. Beersma, R. A. Hut, Gold, silver or bronze: circadian variation strongly affects performance in Olympic athletes. Sci Rep 10, 16088 (2020).

36. N. Zarrouk et al., Time of day effects on repeated sprint ability. Int J Sports Med 33, 975–980 (2012).

37. M. Kjaer, Hepatic glucose production during exercise. Adv Exp Med Biol 441, 117–127 (1998).

38. W. Huang, K. M. Ramsey, B. Marcheva, J. Bass, Circadian rhythms, sleep, and metabolism. J Clin Invest 121, 2133–2141 (2011).

39. A. Lang et al., SIRT4 interacts with OPA1 and regulates mitochondrial quality control and mitophagy. Aging (Albany NY) 9, 2163–2189 (2017).

40. L. Fu et al., SIRT4 inhibits malignancy progression of NSCLCs, through mitochondrial dynamics mediated by the ERK-Drp1 pathway. Oncogene 36, 2724–2736 (2017).

41. Y. Minegishi, A. Otsuka, N. Ota, K. Ishii, A. Shimotoyodome, Combined Supplementation of Pre-Exercise Carbohydrate, Alanine, and Proline and Continuous Intake of Green Tea Catechins Effectively Boost Endurance Performance in Mice. Nutrients 10 (2018).

42. A. Walvekar, Z. Rashida, H. Maddali, S. Laxman, A versatile LC-MS/MS approach for comprehensive, quantitative analysis of central metabolic pathways. Wellcome Open Res 3, 122 (2018).

