## Supplementary figures & material for "Diurnal variation in skeletal muscle mitochondrial function dictates time of day-dependent differences in exercise capacity"

**Supplementary figure legends:**

Figure S1: Time of the day-dependent differences in exercise adaptations, related to figure 1: (A) Total run time of animals at four circadian time points: ZT3, ZT9, ZT15, and ZT21. Data are presented as mean ± SD (n=10). (B-D) Indirect calorimetry measurements during exercise at ZT3 and ZT15: (B) Respiratory Exchange Ratio (RER), (C) Carbohydrate oxidation, and (D) Fat oxidation. Data are presented as mean ± SD (n=5-7). (E) Volcano plot representing metabolites significantly altered in ZT15 sedentary animals compared to ZT3 sedentary animals. The dotted horizontal and vertical lines indicate significance thresholds (p < 0.05 and fold change > 1.5, respectively). Names of significant metabolites are indicated (n=3). (F) Heatmap representation of metabolites altered by exercise at ZT3 and ZT15 relative to corresponding sedentary animals. The scale represents the log2 fold change (log2FC) relative to sedentary animals (n=3). Student's *t*-test (A-E); **p* < 0.05, ***p* < 0.01, ****p* < 0.001, nonsignificant (ns) *p* > 0.05.

Figure S2: time of the-day dependent changes in mitochondrial functions, related to figure 2: (A-B) Quantification of State 3 and State U respiration for both (A) Pyruvate + Malate and (B) Succinate + Rotenone, based on OCR time graphs shown in Figures 2B and 2C, respectively. (C) Quantification of electron flow through Complex I (CI+III+IV), Complex II (CII+III+IV), and Complex IV (CIV), based on OCR time graphs shown in Figure 2D. (D-G) Quantification of mitochondrial morphology from TEM images shown in Figure 2J: (D) Length, (E) Width, (F) Area, and (G) Aspect ratio. Data are represented as violin plots, where the solid horizontal black line indicates the median value, and the dotted lines represent the 1st and 3rd quartiles (n=120-180 mitochondria from 4-5 animals each at ZT3 and ZT15). (H) Quantitative real-time PCR analysis of *Mfn1* mRNA levels in gastrocnemius muscle at ZT3 and ZT15 (n=6). Data are plotted as mean ± SD. Statistical significance was estimated using Student's t-test; *p < 0.05, **p < 0.01, ***p < 0.001, nonsignificant (ns) p > 0.05.

Figure S3: Effect of Sirt4 loss on skeletal muscle physiology, related to figure 3: (A-B) Confirmation of $Sirt4^{-}/^{-}$ genotype: (A) Representative image of genotyping PCR products run on an agarose gel. The expected wild-type (WT) band is 592 bp, and the $Sirt4^{-/-}$ band is 350 bp. (B) Quantitative real-time PCR analysis of Sirt4 gene expression in gastrocnemius muscle from WT and $Sirt4^{-/-}$ mice (n=3). (C-F) Assessment of skeletal muscle atrophy in gastrocnemius muscle from WT and $Sirt4^{-/-}$ mice: (C) Representative images of Haematoxylin-Eosin staining of gastrocnemius muscle sections. (D) Representative images of gastrocnemius muscles, with a scale bar used for qualitative assessment of tissue aspect ratio (N=2, n=3). (E) Bar graph showing the weight of gastrocnemius muscle normalized to body weight (B.W) of the corresponding animal (n=8). (F) Quantitative real-time PCR analysis of *Fbxo32* (Atrogin1) and *Trim63* (Murf1) mRNA levels, normalized to (*Actb*)Actin b, with fold changes relative to WT plotted in a bar graph (N=3, n=3). (G) Quantitative real-time PCR analysis of *Myhc7*, *Myhc2*, and *Myhc6* mRNAs, normalized to Actin b mRNA (N=3, n=3). (H-I) Assessment of glycogen metabolism in gastrocnemius muscle from WT and $Sirt4^{-/-}$ mice: (H) Bar graph showing fold change in normalized glycogen content in $Sirt4^{-/-}$ compared to WT (n=11-12). (I) Quantitative real-time PCR analysis of glycogen metabolism enzymes in gastrocnemius muscle, with fold changes relative to WT plotted in a bar graph (n=4). (J) Quantitative real-time PCR analysis of core clock genes in WT and $Sirt4^{-/-}$ gastrocnemius muscle (n=9). Data in panels B, E-J are presented as mean ± SD. Statistical significance was assessed using Student's t-test (B, E-I) or two-way ANOVA (J); *$p < 0.05$, **$p < 0.01$, ***$p < 0.001$, nonsignificant (ns) $p > 0.05$.

Figure S4: Time of the day-dependent changes in mitochondrial functions in $Sirt4^{-/-}$ skeletal muscle, related to figure 4: (A-D) Quantification of mitochondrial morphology from TEM images shown in Figure 2J: (A) Length, (B) Width, (C) Area, and (D) Aspect ratio of mitochondria from $Sirt4^{-/-}$ animals at ZT3 and ZT15. Data are represented as violin plots, where the solid horizontal black line indicates the median value, and the dotted lines represent the 1st and 3rd quartiles (n=130-160 mitochondria from $Sirt4^{-/-}$ animals at each time point). (E) Quantitative real-time PCR analysis of Mfn1 mRNA levels in gastrocnemius muscle from $Sirt4^{-/-}$ animals at ZT3 and ZT15 (n=6). Data for WT at ZT3 and ZT15 from Figure S2H are used for comparison. Fold changes are calculated relative to WT ZT3 and are presented in bar graphs. All data are plotted as mean ± SD. Statistical significance was assessed using Student's t-test or two-way ANOVA; *p < 0.05, **p < 0.01, ***p < 0.001, nonsignificant (ns) p > 0.05.

Figure S5: The effect of loss of Sirt4 on time of the day dependent-exercise adaptations, related to figure 5: (A-C) Indirect calorimetry measurements during exercise in Sirt4$^{-/-}$ animals at ZT3 and ZT15: (A) RER, (B) Carbohydrate oxidation, and (C) Fat oxidation. Data are presented as mean ± SD (n=5-7). (D) Heatmap of metabolite alterations in response to exercise at ZT3 and ZT15, relative to corresponding Sirt4$^{-/-}$ sedentary animals. The scale represents the log2 fold change (log2FC) compared to sedentary animals (n=3). Statistical significance was assessed using Student's t-test; *p < 0.05, **p < 0.01, ***p < 0.001, nonsignificant (ns) p > 0.05.

**Table 1: Metabolites detection from mouse gastrocnemius muscle**

| Metabolites | Q1 (Parent) | Q3 (Product) | CE | RT |
|---|---|---|---|---|
| Amino acid (Positive polarity mode) | | | | |
| Histidine (His) | 156.0 | 110.0 | 12 | 2.76 |
| Phenylalanine (Phe) | 166.0 | 103.0 | 25 | 6.90 |
| Tyrosine (Tyr) | 182.2 | 136.0 | 14 | 7.04 |
| Tryptophan (Trp) | 205.0 | 146.0 | 15 | 10.0 |
| Alanine (Ala) | 90.0 | 44.0 | 11 | 3.30 |
| Serine (Ser) | 106.0 | 60.0 | 13 | 3.27 |
| Valine (Val) | 118.0 | 55.0 | 15 | 4.35 |
| Leucine (Leu) | 132.0 | 86.0 | 11 | 6.90 |
| Isoleucine (IIe) | 132.0 | 86.0 | 11 | 6.53 |
| Glutamate (Glu) | 148.1 | 84.1 | 16 | 3.48 |
| Glutamine (Gln) | 147.0 | 84.0 | 16 | 3.36 |
| Aspartate (Asp) | 134.1 | 74.0 | 15 | 3.46 |
| Asparagine (Asn) | 133.0 | 74.0 | 15 | 3.30 |
| Threonine (Thr) | 120.0 | 57.0 | 30 | 3.39 |
| Arginine (Arg) | 175.2 | 60.2 | 14 | 2.79 |
| Proline (Pro) | 116.0 | 70.0 | 11 | 3.84 |
| Lysine (Lys) | 147.0 | 84.0 | 16 | 2.72 |
| SAM | 399.0 | 250.0 | 13 | 2.86 |
| SAH | 385.0 | 136.0 | 19 | 6.92 |
| GSH | 308.0 | 162.0 | 17 | 3.08 |
| GSSG | 613.0 | 231.0 | 34 | 7.45 |
| Cystathionine | 223.0 | 134.0 | 14 | 3.19 |
| Homocysteine | 136.0 | 90.0 | 15 | 4.07 |
| Methionine(Met) | 150.1 | 56.0 | 15 | 5.30 |
| 13C4-Aspartate | 138.9 | 91.1 | 15 | 3.46 |
| Nucleotide (Positive polarity mode) | | | | |
| AMP | 348.2 | 136.0 | 21 | 6.62 |
| GMP | 364.2 | 152.1 | 18 | 7.96 |
| CMP | 324.2 | 112.1 | 21 | 4.57 |
| UMP | 325.2 | 113.1 | 21 | 7.83 |
| Sugar Phosphate metabolites (Negative polarity mode) | | | | |
| UDP-Glc | 565.0 | 323.0 | -25 | 3.35 |
| UDP-GlcNAc | 605.8 | 384.9 | -37 | 3.37 |
| G3P | 169.0 | 97.0 | -20 | 3.30 |
| 3PG | 185.0 | 97.0 | -20 | 3.35 |
| G6P | 259.0 | 97.0 | -20 | 3.39 |
| 6PG | 275.0 | 97.0 | -20 | 3.40 |
| R5P | 229.0 | 97.0 | -20 | 3.36 |
| S7P | 289.0 | 97.0 | -20 | 3.34 |
| 13C4-Aspartate | 138.9 | 91.1 | -17 | 3.46 |
| TCA metabolite derivatization (Positive polarity mode) | | | | |
| MG | 283.0 | 91.2 | 25 | 11.3 |
| Lactate | 196.0 | 65.2 | 40 | 6.71 |
| Pyruvate | 299.0 | 181.0 | 15 | 9.37 |

| | | | | |
|---|---|---|---|---|
| Oxaloacetate | 448.0 | 91.2 | 25 | 9.12 |
| Citrate | 508.0 | 91.2 | 25 | 8.36 |
| 2-Keto Glutarate | 462.0 | 339.0 | 11 | 9.29 |
| Fumarate | 327.0 | 204.0 | 15 | 8.10 |
| Succinate | 329.0 | 206.0 | 15 | 7.80 |

### Table 2: List of primers used for real-time qPCR

| Gene name | Primer | Sequence (5' to 3') |
|---|---|---|
| ***Myhc7*** | FP | AGCATTCTCCTGCTGTTTCC |
| ***Myhc7*** | RP | ACTCTTCTTTGTCATCGGGC |
| ***Myhc6*** | FP | GAAGTTGCATCCCTAAAGGCAG |
| ***Myhc6*** | RP | CAAAAGGCTTGTTCTGAGCC |
| ***Myhc2*** | FP | TCACCTACCAGACCGAGGAG |
| ***Myhc2*** | RP | ACCTCTCAACAGAAAGATGGA |
| ***Gys1*** | FP | AAGGTGACAGGGGATGAATG |
| ***Gys1*** | RP | ACGCCCAAAATACACCTTAC |
| ***Pygm*** | FP | TAGGTTTATGGTGCCGAGG |
| ***Pygm*** | RP | CGGCGGGAATAACTTTCTC |
| ***Gbe1*** | FP | TGCTCTATCATCACCACGG |
| ***Gbe1*** | RP | ATCCCTGATACATCCTCTGC |
| ***Agl1*** | FP | ACATGAAGGACGAGGGTTTC |
| ***Agl1*** | RP | TTCCAGCAACCAACGAACAG |
| ***Fbx32*** | FP | ACGTAGTAAGGCTGTTGGAG |
| ***Fbx32*** | RP | ACCAGTGTGCATAAGGATGT |
| ***Trim63*** | FP | GGCTACCTTCCTCTCAAGTG |
| ***Trim 63*** | RP | CTTCTTTACCCTCTGTGGTCAC |
| ***Mfn1*** | FP | CAGGGACGGAGTGAGTGTC |
| ***Mfn1*** | RP | GCTTCAGTGGAGATACCGTT |
| ***Actb*** | FP | CCTCCCTGGAGAAGAGCTATGA |
| ***Actb*** | RP | GCACTGTGTTGGCATAGAGGTC |
| ***18S rRNA*** | FP | TTTCGAGGCCCTGTAATTGG |
| ***18s rRNA*** | RP | CCCAAGATCCAACTACGAGC |
| ***Clock*** | FP | TTGCTCCACGGGAATCCTTC |
| ***Clock*** | RP | CTACTGTGGCTGGACCTTGG |
| ***Arntl*** | FP | AATGAGCCAGACAACGAGGG |
| ***Arntl*** | RP | GCTGTCGCCCTCTGATCTAC |
| ***Cry1*** | FP | CACTGGTTCCGAAAGGGACTC |
| ***Cry1*** | RP | CTGAAGCAAAAATCGCCACCT |
| ***Per2*** | FP | CTCCAGCGGAAACGAGAACT |
| ***Per2*** | RP | CTCACTACTGCAGCCGCTC |

**Table 3: List of antibodies used for western blot**

| Antibody name | Company | Catalog number | Dilution |
|---|---|---|---|
| **Mouse IgG-HRP** | Sigma | A9044 | 1:8000 |
| **Rabbit IgG-HRP** | Sigma | A0545 | 1:8000 |
| **Actin** | Sigma | A1978 | 1:8000 |
| **Tubulin** | Sigma | T8328 | 1:8000 |
| **Atrogin 1** | Abcam | ab168372 | 1:1000 |
| **TOM20** | Cell Signaling | 42406T | 1:1000 |
| **PGC1a** | Invitrogen | PA5-38021 | 1:1000 |
| **VDAC1** | Abcam | ab15895 | 1:1000 |
| **Total OXPHOS antibody** | Abcam | ab110413 | 1:1000 |
